# Dysregulated dsRNA sensor signaling and viral infection during onset of pediatric autoimmune interferonopathy

**DOI:** 10.64898/2026.05.27.728148

**Authors:** Thomas RJ Moreau, Yann Aquino, Yixiang YJ Zhu, Vincent Bondet, Chloé Albert-Vega, Françoise Donnadieu, Florian Dubois, Baptiste Periou, Farah Rahal, Mina Tizerarine, Saskia R Veldkamp, Etienne Villain, Anthony Bertrand, Christine Bodemer, Camille Brunaud, Marie-Louise Frémond, Benjamin Fournier, Bénédicte Hoareau, Pierre Quartier, François-Jérôme Authier, Eugénie Sarda, Adrien Schvartz, Angélique Vinit, Annet van Royen-Kerkhof, Femke van Wijk, Anne Welfringer-Morin, Frédéric Rieux-Laucat, Marc Jansen, David Hing, Tatiana Traboulsi, Carolina Moraes Cabé, Milena Hasan, David Hardy, Michael White, Lluis Quintana-Murci, Isabelle Melki, Brigitte Bader-Meunier, Cyril Gitiaux, Mathieu P Rodero, Darragh Duffy

## Abstract

Juvenile dermatomyositis (JDM) is characterized by a type I interferon (IFN-I) signature associated with disease activity. We previously identified a link between SARS-CoV-2 infection and the onset or relapse of JDM. Here, we show that newly diagnosed JDM patients display an overexpression of *IFIH1* (encoding MDA5 protein) at baseline, coupled with an altered response to dsRNA stimulation at proteomic and transcriptomic levels, indicating abnormal activation of this antiviral sensing pathway. Single-cell transcriptomic and chromatin accessibility profiling of peripheral blood mononuclear cells (PBMCs) further revealed myeloid-specific enrichment of interferon-stimulated genes (ISGs) and preferential disruption of this pathway at disease onset, supporting a dysregulated IFN-I state in this cell type. We identified SARS-CoV-2 RNA in muscle biopsies of two Covid-19 pandemic-onset JDM patients, strongly implicating viral infection as a potential trigger of the dysregulated MDA5 immune response. To extend these observations beyond SARS-CoV-2, we screened two independent retrospective cohorts for antibodies against 27 common childhood infections. In our discovery cohort JDM patients showed significantly increased exposure to 4 RNA viruses in line with our immunological findings. Increased exposure to RSV B was confirmed in an independent replication cohort supporting a robust association with JDM pathophysiology. Together, these findings integrate systemic, single-cell, and tissue-level analyses implicating RNA viral infection and biased antiviral sensing in shaping IFN-I responses at JDM onset, providing mechanistic insight into environmentally triggered pathogenesis.

**One sentence summary:** Type I interferon dysregulation at juvenile dermatomyositis onset implicates altered dsRNA sensing and RNA viral exposure as potential disease triggers.

## INTRODUCTION

Juvenile dermatomyositis (JDM) is an inflammatory, autoimmune, pediatric-onset disease. A reported incidence of 1.6 to 4 cases per million children per year with significant morbidity and mortality (2%) underlines both the rarity and the severity of this disease [^1,2,3,4,5^]. The main symptoms of JDM include characteristic skin conditions such as heliotrope rash or Gottron’s papules and symmetric proximal muscle weakness [^2^]. Corticosteroids (CTC) combined with methotrexate (MTX) remain the leading therapeutic approach for managing these inflammatory symptoms, with studies demonstrating their efficacy in reducing mortality [^3,6,7^]. However, these treatments often lack efficacy [^6^], and present a risk of resistance and adverse effects such as infection and growth retardation. Recently, treatments targeting the IFN-I pathway, including JAK inhibitors and antibodies against IFN-β and IFNAR, have shown promising results in dermatomyositis (DM), but response rates remain highly variable [^8,9,10,11,12^]. Thus, it is essential to better understand the pathophysiological mechanisms underlying JDM to develop personalized treatment strategies.

JDM is characterized by a type I interferon (IFN-I) signature [^13^], with interferon-stimulated genes (ISGs) upregulated in both blood and affected tissues [^14,15,16,17^], which is associated with disease severity [^18,19^]. IFN-I production is most commonly induced after viral infection following sensing by pattern recognition receptors (PRR) such as Toll-like receptors (TLRs), RIG-I like receptors (RLRs) or intracellular DNA sensors like cGAS. The secreted IFN-I binds to the IFNAR receptor resulting in activation of hundreds of ISGs. Recent studies have highlighted the potential of environmental factors such as viral infections in triggering autoimmune diseases, several of which are characterized by an IFN-I signature [^20,21,22^].

In JDM, several case reports have described a temporal association between various infectious pathogens and disease onset [^23^]. Additionally, we recently showed that among 10 new-onset or relapsed cases of JDM, two had a concurrent history of SARS-CoV-2 infection [^24^]. However, the biological pathways linking pathogen exposure to disease onset or relapse have not yet been elucidated. We hypothesized that dysregulation of specific viral nucleic acid sensor pathways may be implicated in JDM. To test this hypothesis, we performed a systems immunology study on a prospective cohort of JDM patients recruited at disease onset (**Fig.1**). We observed a striking defect in double-stranded RNA (dsRNA) sensing pathway in JDM patients, accompanied by increased serological evidence of RNA virus exposure and direct detection of SARS-CoV-2 nucleocapsid (NP) RNA in skeletal muscle biopsies, supporting a viral contribution to disease pathogenesis.

**Figure 1.**
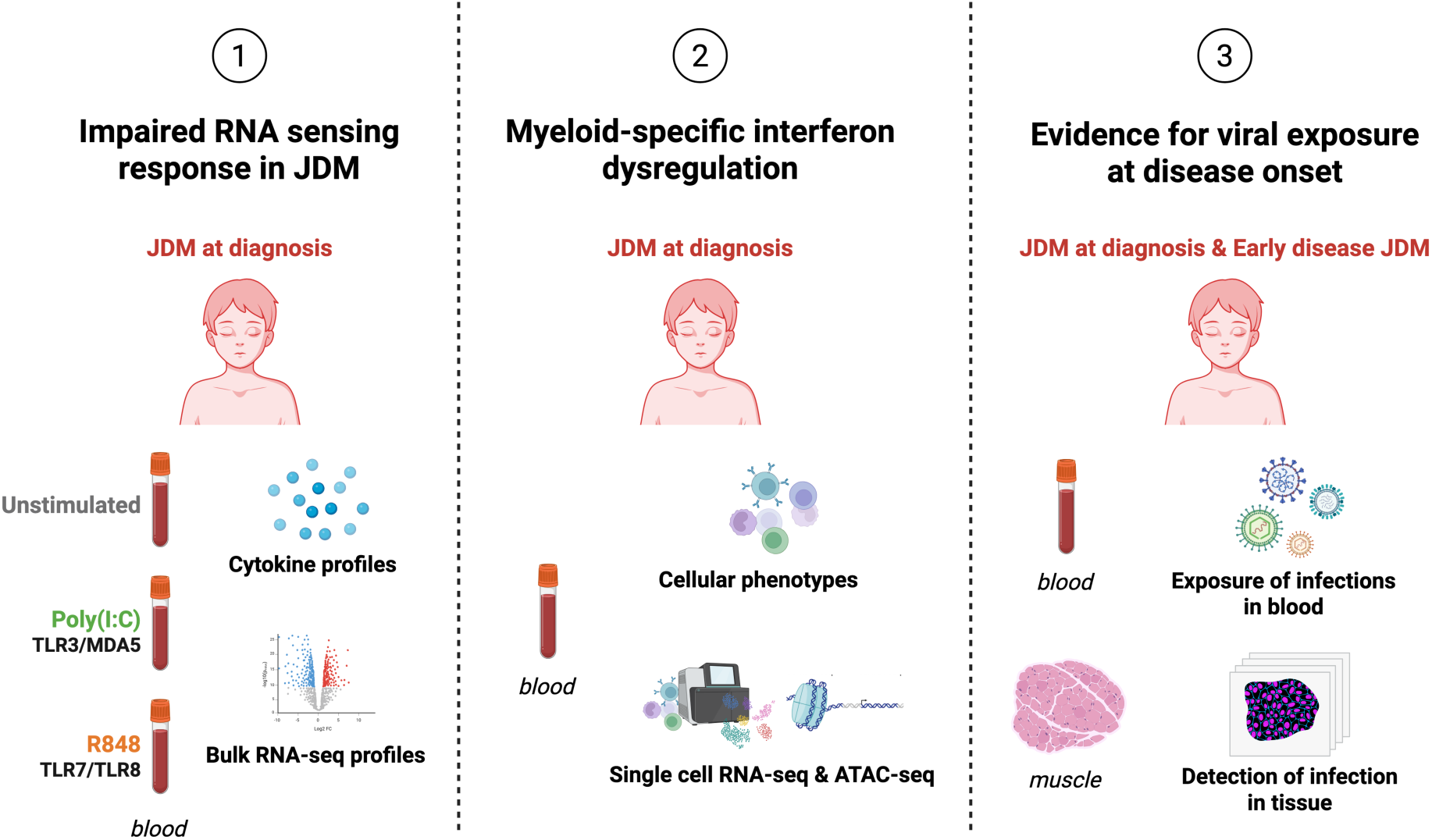
Overview of the experimental design.

## RESULTS

### JDM patients show cytokine and transcriptome IFN-I signature at diagnosis

We first investigated whether JDM patients, at diagnosis and in the absence of prior treatment, had a specific systemic cytokine profile. Of the 46 cytokines measured by Luminex and Simoa digital ELISA in TruCulture supernatants, IFN-α, IFN-β, CXCL10, IL-10, CCL11, CCL2, CCL4, and IL-1ra all showed elevated circulating levels in JDM patients at diagnosis (n=11) compared to controls (n=10) (**Fig.2A**). All JDM patients exhibited a significant IFN-I signature (IFN-α, IFN-β, and ISGs: CXCL10, CCL2), along with significantly elevated levels of anti-inflammatory cytokines IL-10 and IL-1ra (**Fig.2B**). These results are consistent with the previously reported IFN-I signature in DM patients at different stages of the disease [^25,26,27,28^]. We then performed RNA-seq analysis to confirm these results at the transcriptomic level. The most differentially expressed upregulated genes (DEGs) were known ISGs or genes related to the IFN-I pathway (**Fig.2C**). In JDM, 324 Gene Ontology (GO) pathways were upregulated compared to controls, primarily involving viral response, immune pathways, and IFN-I related pathways (**Fig.2D, Table S1**). Neutrophil-enriched pathways were also observed (**Table S1**), confirming previous results reporting a neutrophil signature associated with disease activity in JDM [^29^]. Five downregulated pathways were identified in newly diagnosed JDM, primarily related to T cells, including downregulation of the *CD8A* gene. A reduced peripheral blood CD8⁺ T cell count has previously been reported in treatment-naïve dermatomyositis patients [^30^] (**Table S2**). Overall, these findings confirm a robust IFN-I signature in JDM at diagnosis at both proteomic and transcriptomic levels.

**Figure 2.**
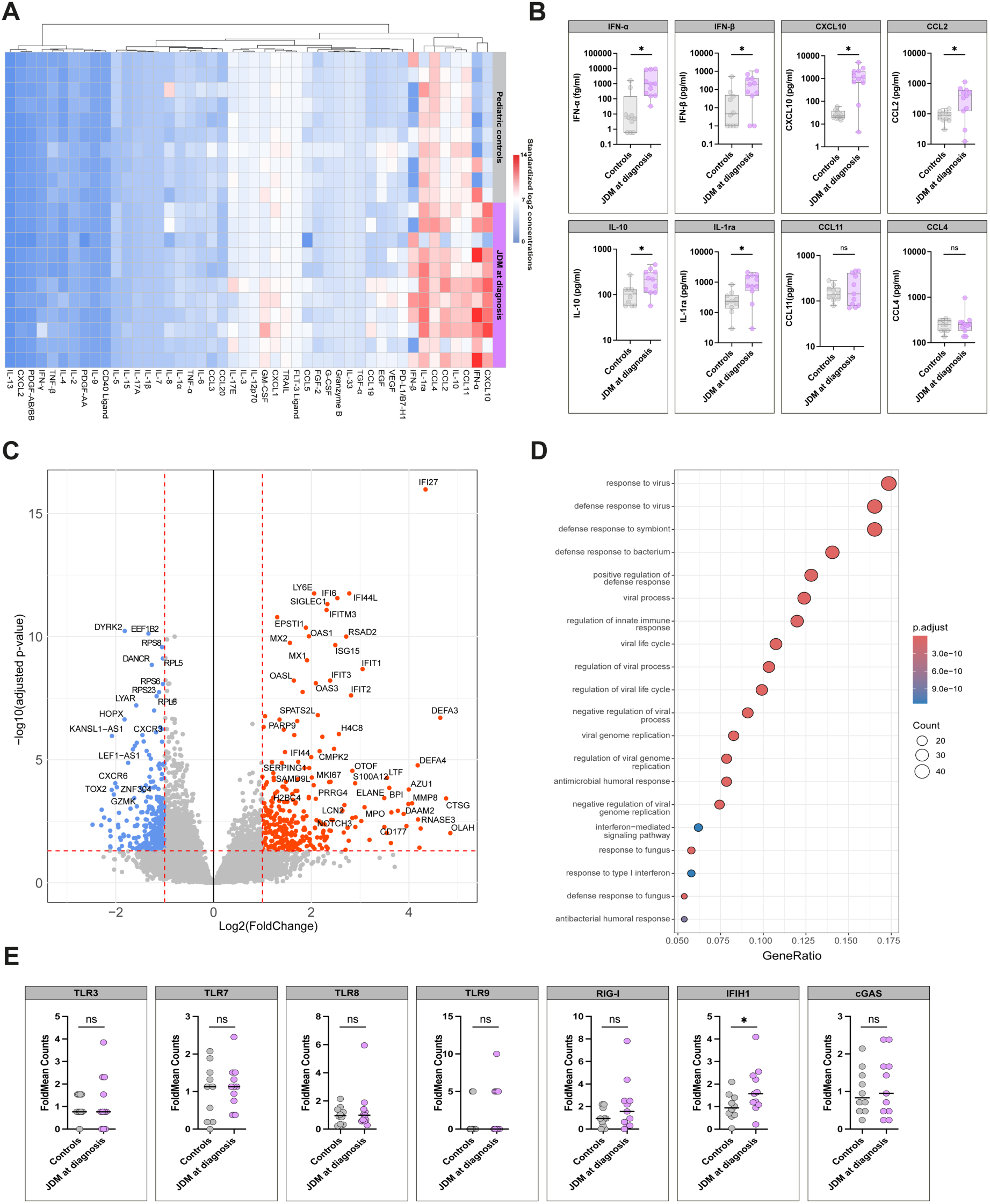
Proteomic and transcriptomic analysis of JDM patients at diagnosis shows a strong IFN-I signature. **(A)** Heatmap of standardized log2 concentrations for 46 cytokines in 11 JDM patients at diagnosis versus 10 age-matched pediatric controls. Cytokine columns are ranked using a hierarchical clustering method. **(B)** Boxplots showing the 8 cytokines from the cluster identified in panel A that displayed distinct profiles in JDM patients at diagnosis compared with controls. Each dot represents an individual patient or control, and the line indicates the median. **(C)** Volcano plot showing differential gene expression in JDM patients at diagnosis compared to controls. Genes with p-adj < 0.05 and |log2FC| ≥ 1 are highlighted (upregulated in red, downregulated in blue), while genes not meeting these thresholds are shown in grey. **(D)** Enriched pathways in JDM at diagnosis compared to controls. The count represents the number of genes from DEGs that are found in a specific Gene Ontology (GO) term. The GeneRatio is the proportion of our DEGs associated with a particular GO term relative to the total number of our DEGs. **(E)** Scatter plots of FoldMean counts for specific PRR genes in JDM at diagnosis versus controls, with lines indicating medians. FoldMean represents each patient’s or control’s count relative to the mean control count for each gene. Mann-Whitney test was performed for statistical comparisons, p-values were corrected using False Discovery Rate (FDR) method. * p-adj < 0.05. (n=11 JDM, 10 controls).

As several studies have reported that the onset of JDM was associated with viral infections [^23^], we hypothesized that the strong IFN-I signature in JDM may be driven by dysregulated PRR signaling. Therefore, we examined the expression levels of various PRRs including *TLR3*, *TLR7* and *TLR8* that recognize ssRNA viruses, *TLR9* that recognizes DNA viruses, RLRs (*RIG-I* and *IFIH1*, encoding MDA5, which detect cytosolic viral dsRNA), as well as the intracellular DNA sensor *cGAS* (**Fig.2E**). Interestingly, *IFIH1* was the only viral sensor gene over-expressed in JDM patients at diagnosis compared to controls. The expression levels of *IFIH1* was correlated with *RSAD2* and *IFIT1*, two ISGs that are part of an ISG signature identified in several pediatric inflammatory diseases (**Fig.S1**) [^31^]. These results suggest that *IFIH1/*MDA5 may be implicated in the IFN-I signature in newly diagnosed JDM. Interestingly, autoantibodies (aab) targeting the MDA5 protein define a subgroup of JDM patients characterized by severe cutaneous manifestations and pulmonary complications, though only 3/11 of the onset patients in our study were anti-MDA5 aab positive (**Table S3**) [^32^].

### JDM patients exhibit a defective cytokine response to Poly(I:C) stimulation at diagnosis

To test for functional relevance of this *IFIH1* overexpression in JDM patients at diagnosis we stimulated patient and control blood with Poly(I:C), which mimics dsRNA viruses and activates TLR3 and MDA5 pathways. As a positive control, we used R848 which mirrors single-stranded RNA (ssRNA) viruses and binds to TLR7 and TLR8 (**Table S4**). Following Poly(I:C) stimulation, controls showed a significant induction of 12 cytokines related to antiviral and innate immune responses (**Fig.3A, Table S4**). Strikingly, IFN-α and the downstream chemokines CXCL10 and CCL2 were not induced in JDM patients (**Fig.3A**). Among the 46 cytokines measured, only IFN-β and IL-2 were significantly, albeit weakly, increased in JDM patients (**Table S4**). In contrast after stimulation with the TLR7/8 agonist R848, both controls and JDM patients significantly produced IFN-I, ISGs and other cytokines involved in antiviral and immune responses compared to their basal state (Null). Cytokine responses to R848 were comparable between JDM patients and controls despite higher baseline levels in JDM. Therefore, the defective response in newly diagnosed JDM patients appears to be specific to Poly(I:C) stimulation, implying a dysregulation of dsRNA sensing at disease onset. These findings coupled with the observed *IFIH1* overexpression at baseline suggests that MDA5 may be specifically implicated in the defective dsRNA response.

**Figure 3.**
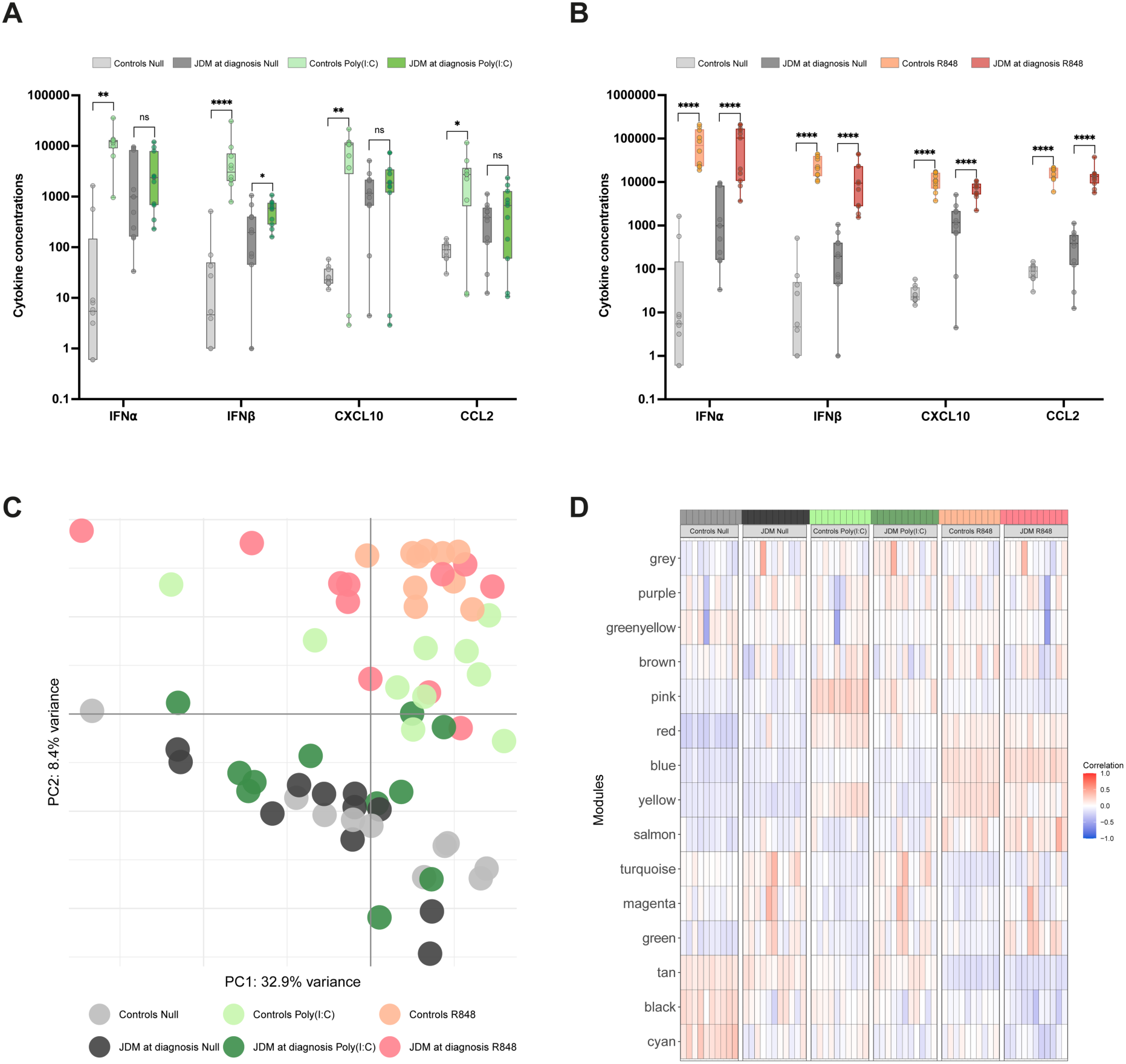
Defective proteomic and transcriptomic response to Poly(I:C) stimulation in JDM patients at diagnosis. **(A–B)** Cytokine concentrations (IFN-α, IFN-β, CXCL10 and CCL2) measured in the supernatant of TruCulture supernatants containing whole blood from JDM patients at diagnosis and healthy controls. (A) Poly(I:C) stimulation. (B) R848 stimulation. Unstimulated conditions (Null) are shown for JDM patients (dark grey) and controls (light grey). Stimulated conditions are shown for JDM patients (Poly(I:C): dark green; R848: red) and controls (Poly(I:C): light green; R848: orange). Statistical significance between JDM patients and controls was assessed using a Kruskal–Wallis test followed by Dunn’s multiple comparisons test. * p < 0.05, ** p < 0.01, *** p < 0.001, **** p < 0.0001. **(C)** Principal component analysis (PC1 and PC2) of gene expression profiles obtained by bulk RNA-seq from JDM patients at diagnosis and controls, under unstimulated conditions (Null) or after Poly(I:C) or R848 stimulation. **(D)** Heatmap displaying correlations between gene co-expression modules at baseline/Poly(I:C)/R848 condition. Color intensity reflects the correlation between genes module and each condition. (n=11 JDM, 10 controls).

### Transcriptomic analysis confirms the defective response to Poly(I:C) in JDM at diagnosis

We next performed transcriptomic analyses to explore the underlying factors contributing to dysregulated dsRNA sensing in JDM at diagnosis. Principal component analysis (PCA) of the Null, Poly(I:C) and R848 samples showed that stimulation and disease status explain most of the variance between samples, as captured by the first two principal components of the data (41.3%) (**Fig.3C**). R848 stimulated samples from JDM (in red) and controls (in orange) clustered together (on the top right) and are separated from Null samples (on the bottom), indicating important differences after stimulation. Control samples after Poly(I:C) stimulation (in light green) also clustered separately from Null samples (in grey) and R848 samples. In contrast, and in accordance with previous proteomic findings, 9 out of 11 Poly(I:C) samples from JDM patients (dark green) cluster together with the Null condition.

We conducted DEG analysis to identify specific genes that may be implicated in the dysregulation of the TLR3/MDA5 pathway in newly diagnosed JDM patients. The number of DEGs varied following TLR stimulations (**Fig.S2**). After Poly(I:C) stimulation, controls exhibited 1580 DEGs, whereas JDM patients showed only 71 DEGs compared to their respective baseline (Null). This shows that the transcriptomic profile of JDM patients following Poly(I:C) stimulation is highly similar to their baseline transcriptomic profile. In contrast, R848 stimulation resulted in a substantial number of DEGs in both JDM and controls (respectively 1616 and 2567 DEGs). To evaluate similarities in the stimulated responses between controls and JDM patients, we examined the overlap of the up- or down-regulated DEGs between both groups. Of the 71 DEGs detected after Poly(I:C) stimulation in JDM, 3 upregulated genes and 7 downregulated genes were observed exclusively in JDM patients. (**Fig.S2**). As expected, no pathways were enriched in JDM patients after Poly(I:C) stimulation. In contrast, after R848 stimulation, we observed enriched pathways in both JDM and controls, the majority of them related to antiviral responses, innate immunity signaling and the IFN-I pathway (**Fig.S3**).

Next, we conducted weighted gene co-expression analysis (WGCNA) to identify clusters of genes with similar expression patterns (key gene modules). Following Poly(I:C) and R848 stimulation, 15 gene modules were identified using hierarchical clustering and the dynamic tree cut method (**Fig.S4**). The ‘turquoise’, ‘greenyellow’ and ‘red’ modules showed high associations with Null, Poly(I:C) and R848 conditions in JDM patients but not in controls, suggesting a potential JDM disease signature (**Fig.3D**). Genes in these modules were associated with several processes such as leukocyte migration, neutrophil activation, and reactive oxygen species metabolism, corroborating our observations and previous studies that reported these pathways enriched in JDM patients [^29,31,33^] (**Fig.S5)**. Module pattern associations showed striking similarities between JDM Null and JDM Poly(I:C), indicating a common gene expression in these two conditions. In contrast, for R848 stimulation both JDM and controls had a similar pattern with ‘blue, ‘black’ and ‘brown’ modules having strong associations. These three gene modules were also associated with control Poly(I:C) responses and were implicated in antiviral responses and innate immunity. Overall, we confirmed the defective response to Poly(I:C) in JDM at diagnosis at transcriptomic levels.

### Cellular characterization of JDM at diagnosis

Given the impaired IFN-I response to Poly(I:C) stimulation, we next sought to define a potential cellular basis of this dysregulation in JDM at diagnosis. Mass cytometry phenotyping of the main circulating immune cell populations (**Fig.4A**) showed few significant differences between JDM patients and controls. Specifically, no significant differences were observed in proportions of monocytes or pDCs, the main IFN-I producing cells. We did observe a significant difference (p-adj = 0.026) in basophil proportions, which were lower in JDM patients than controls (**Fig.4B**). An accumulation of basophils in the skin lesions of dermatomyositis patients has previously been reported [^34^] suggesting that lower levels of basophils in the blood of JDM patients may be due to their migration into affected tissues such as skin. Similarly, when examining B cell subsets, we observed a significant (p-adj = 0.026) decreased frequency of central memory B cells (CD19^+^CD27^+^ CD38^lo^) compared to controls (**Fig.4C**). This is consistent with previously reported observations, where an increased presence of memory B cells was found in inflamed muscle tissue, accompanied by a decrease in peripheral blood in a single JDM patient [^35^]. We also observed a significant (p-adj = 0.026) decrease of effector memory CD8 T cells within CD8 T cell subsets (**Fig.4C**). This finding is consistent with our bulk RNA-seq data showing downregulation of CD8⁺ T cell–related signaling pathways in JDM patients. Together with previous reports of increased CD8⁺ memory T cell infiltration in muscle tissue correlating with disease activity [^36^], these data support a redistribution of CD8⁺ T cells toward inflamed tissues in JDM. Overall, using mass cytometry, we did not identify alterations in immune cell subset frequencies that could account for the impaired IFN-I response to dsRNA stimulation.

**Figure 4.**
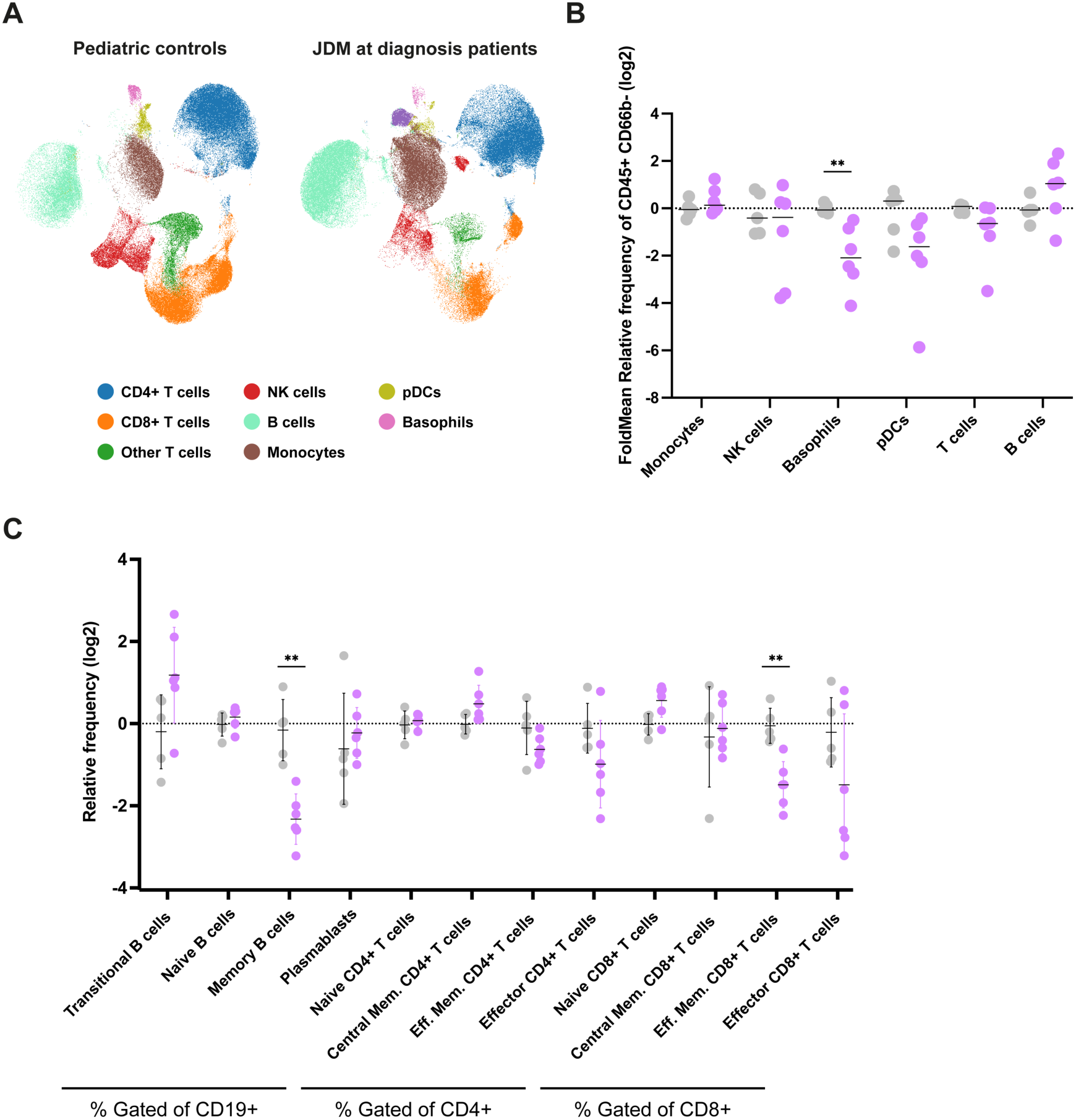
Phenotypic characterization of circulating cells in JDM patients at diagnosis. **(A)** UMAP representation of unsupervised clustering results of immune cell phenotypes of JDM patients at diagnosis and paediatric controls using mass cytometry (CyTOF). **(B)** Manually gated CyTOF cell populations from whole blood of 6 JDM patients at diagnosis versus 5 age-matched paediatric controls. Relative frequency corresponds to the ratio of cell populations of interest / cell populations CD45+ CD66b-. Mann-Whitney test was performed for statistical comparisons, p-values were corrected using False Discovery Rate (FDR) method. Line in each group represents the median. **(C)** Relative frequency of cell populations of interest gated on primary immune phenotypes (CD19+/CD4+/CD8+). Mann-Whitney test was performed for statistical comparisons, p-values were corrected using False Discovery Rate (FDR) method. (n=6 JDM, 5 controls).

### Single-cell RNA sequencing and chromatin accessibility profiling define altered myeloid programs in JDM

Given the lack of differences in the proportions of circulating cells that sense dsRNA to produce IFN-I, we next tested a hypothesis of intra-cellular alterations in immune cell populations. To do this we performed single-cell RNA-sequencing and assays for transposase-accessible-chromatin sequencing (scRNA-seq, scATAC-seq) on peripheral blood mononuclear cells (PBMCs) isolated from a new onset JDM patient and two age matched controls (**Fig.5A**). Consistent with our bulk whole-blood observations at baseline (**Fig.2D**), significant transcriptional changes in the JDM patient (|logFC| > 0.5, FDR < 0.01; **Table S5**) were more than twice as likely to involve IFN-I response genes relative to all other genes expressed in each immune lineage (odds ratio [OR] > 2.4, FDR < 1.4 × 10^-3^). Notably, this enrichment was strongest in myeloid cells (OR = 8.7, FDR < 2.7 × 10^-25^), which also showed the most transcriptional differences between the JDM patient and controls (**Fig.5B**). These findings highlight the myeloid compartment as a key component of JDM aetiology. Among the most significantly enriched pathways in myeloid cells from the JDM patient, upregulated programs were primarily related to signaling and response-to-stimulus processes (**Fig.5C, Table S6**). In contrast, downregulated pathways were biased towards genes involved in protein translation and, strikingly, the immune response, such as *IFI16*, *AIM2* and *NOD2* (FE > 3, FDR < 1.6 × 10^-30^).

**Figure 5.**
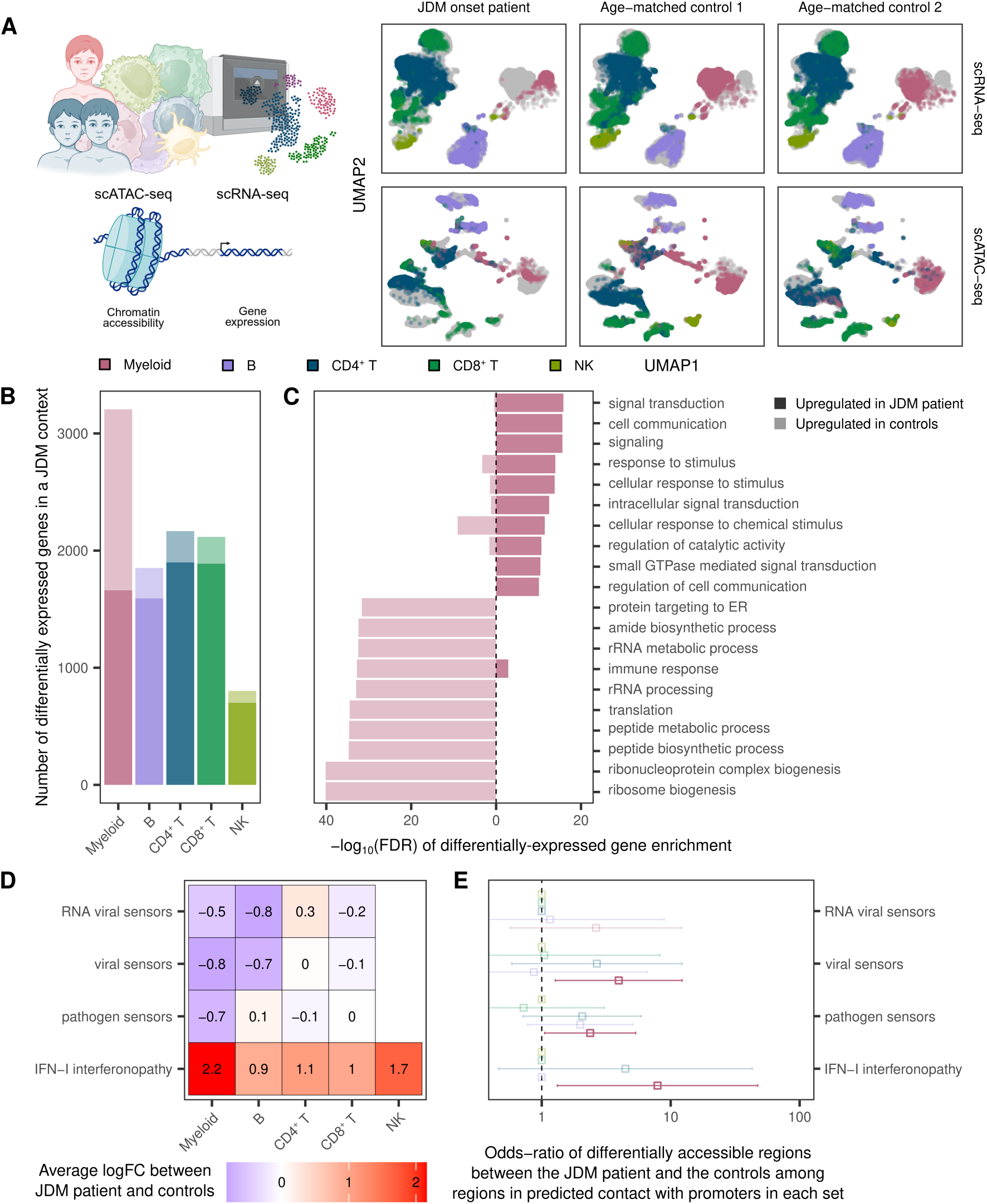
Altered interferon-regulated transcriptional programs in myeloid cells from a JDM patient. **(A)** Single-cell transcriptomic (scRNA-seq) and chromatin-accessibility (scATAC-seq) profiles from peripheral blood mononuclear cells of a JDM onset patient and two age-matched controls projected in UMAP coordinates. Each data point is a single cell or nucleus, colored according to immune lineage identity inferred from transcriptional activity or the accessibility of chromatin around coding genes; the gray background outlines the projection of all cells/nuclei from each donor in each data modality. **(B)** Number of differentially expressed genes (|logFC| > 0.5, FDR < 0.01) in cells from the JDM onset patient relative to the two age-matched controls across immune lineages, stratified by whether they are upregulated in the patient’s (higher opacity) or the controls’ cells. **(C)** Over-representation of differentially expressed genes in myeloid cells from the JDM onset patient among Gene Ontology (GO) biological processes. For the sets of genes upregulated or downregulated relative to the age-matched controls, the ten most significantly enriched GO pathways are shown. **(D)** Effect-size estimates (log-fold change; logFC) of the JDM status averaged across differentially expressed genes in each immune lineage (FDR < 0.05) which belong to each of four custom-built gene sets related to specific anti-viral-defense cellular processes. **(E)** Odds-ratio of differentially accessible open chromatin regions (|log2FC| > 0.5, FDR < 0.01) in each immune lineage among regions linked via activity-by-contact models to the promoters of genes involved in pathogen sensing, as well as type I interferonopathies.

To further understand the transcriptional impact of JDM on the myeloid immune response, we applied a curated IFN-I gene signature previously described in type I interferonopathies including JDM [^31,37,38^] (**Table S7**). We also examined custom gene sets composed of pathogen-sensing receptors (**Table S7**), with a particular focus on viral RNA sensors, given our previous results of defective responses to dsRNA stimulation (**Fig.3**). While the average expression of all pathogen-sensing receptor pathway genes was lower in myeloid and lymphoid cells from the JDM patient, the mean expression of genes associated with type I interferonopathies was nearly five times higher in myeloid cells from the JDM patient relative to controls (average log_2_FC = 2.2, FDR < 0.05; **Fig.5D**). Together these findings suggest a dissociation between upstream antiviral sensing and downstream IFN-I signaling in a JDM context.

Consistent with this hypothesis, analyses of scATAC-seq data revealed that changes in chromatin accessibility were over twice more likely (2.4 < OR < 7.9, *p* < 0.03) to affect putative regulatory regions in predicted contact [^39,40^] with the promoters of pathogen-sensor genes, as well as genes linked to type-I interferonopathies compared to other open chromatin regions genome-wide (**Fig.5E**). Moreover, differentially accessible chromatin regions were most strongly enriched in binding motifs for interferon regulatory factors (FE > 2.6, FDR < 0.01; **Table S8A-B**), specifically in myeloid cells. Overall, our results show epigenetic remodeling of antiviral and interferon-related programs in myeloid cells from a JDM patient, in line with a potential contribution of viral infection to the IFN-I dysregulation observed in JDM.

### Infection by RNA viruses may contribute to IFN-I dysregulation in JDM

Given the altered antiviral sensing and IFN-I signaling programs identified in myeloid cells, we next assessed for evidence of increased viral exposure in JDM patients. Indeed, several reports have documented cases of JDM emerging concomitantly with SARS-CoV-2 infection during the COVID-19 pandemic [^24,41,42^]. Notably, in our center, newly diagnosed JDM cases increased during the pandemic period, and later declined when compared with expected incidence patterns (**Fig.S6**). We quantified IgG levels against SARS-CoV-2 in TruCulture Null supernatants (in the absence of serum) from 13 newly diagnosed JDM patients and 11 age-matched pediatric controls, recruited in parallel during the Covid-19 pandemic. Seven out of the 11 controls were seropositive for SARS-CoV-2 RBD (Receptor binding domain from the Wuhan strain) while all newly diagnosed JDM patients were seropositive (p=0.0311, Fisher contingency test) (**Fig.S7**). Of note, none of the JDM patients had been vaccinated against SARS-CoV-2 at the time of recruitment.

To determine whether viral transcripts were present in diseased tissue, we performed RNAscope in situ hybridization targeting SARS-CoV-2 nucleocapsid (NP) RNA on diagnostic muscle biopsies from two JDM patients diagnosed during the COVID-19 pandemic with documented SARS-CoV-2 infection, and one JDM onset patient prior to the pandemic (**Fig.6A**). In both JDM patients with a PCR confirmed SARS-CoV-2 infection, we detected NP-positive puncta (foci) within the muscle tissue (14 & 24 puncta in either patient). These NP RNA signals appeared spatially clustered within localized regions of the muscle tissue. In contrast, a pre-pandemic JDM control biopsy showed only 3 non clustered NP-positive puncta under identical imaging conditions (**Fig.6B**). This focal enrichment of SARS-CoV-2 NP RNA indicates viral replication in the muscle of patients at disease onset.

**Figure 6.**
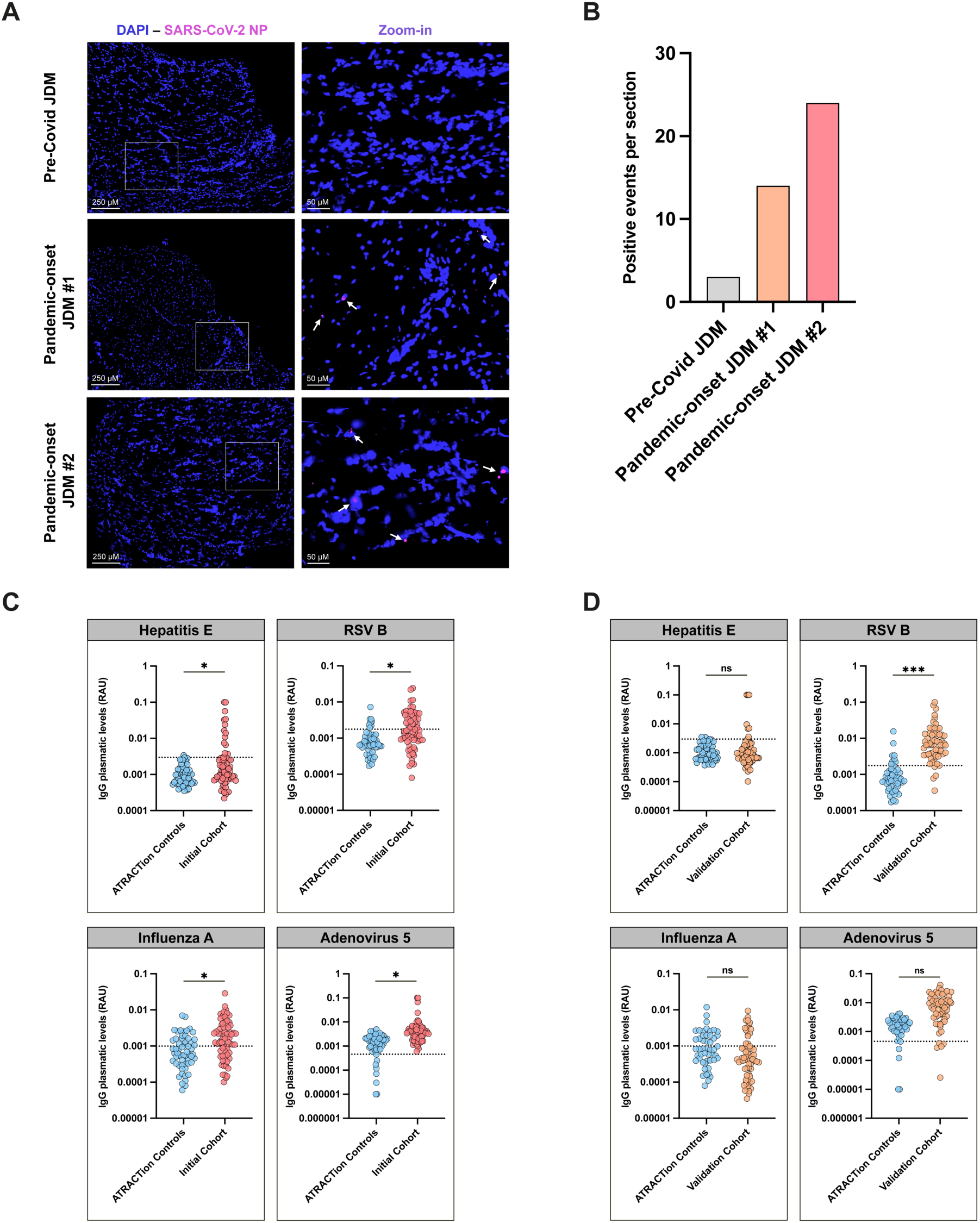
Association between viral infection and JDM. **(A)** Representative RNAscope fluorescence images showing detection of SARS-CoV-2 nucleocapsid (NP) RNA in muscle biopsies from JDM patient recruited before Covid-19 pandemic (pre-Covid JDM) and 2 JDM patients at diagnosis with a suspicion of Covid infection. RNAscope puncta corresponding to viral RNA (SARS-CoV-2 NP) are shown in pink. Nuclei are stained with DAPI (blue). **(B)** Quantification of SARS-CoV-2 NP RNA RNAscope puncta across entire muscle sections. Each point represents a detected RNAscope signal event quantified using QuPath analysis. **(C)** Out of the 27 viral antigens tested, JDM patients showed significantly higher seropositivity compared to age-matched pediatric controls to Hepatitis E, RSV B, Influenza A, and Adenovirus 5. **(D)** Among the significant antigens identified in our initial cohort, JDM patients in the validation cohort shows increased seropositivity specifically against RSV B.

To explore the potential involvement of other viruses to JDM onset we then extended our analysis to a larger retrospective cohort, including patients recruited outside of the pandemic. We applied multiplexed IgG Luminex assays on plasma samples from 68 JDM patients from France (initial cohort) recruited from 2017 to 2023. We quantified IgG levels against 27 different microbial pathogens (including DNA and RNA viruses, parasites and bacteria listed in **Table S9**). This JDM cohort had a significantly higher proportion of patients seropositive for 4 viruses compared to age-matched controls (ATRACTion healthy control cohort): hepatitis E, respiratory syncytial virus group B (RSV B), Influenza A and adenovirus type 5 (ADE5) (**Fig.6C, Table S10**). To confirm these results, we assessed plasma IgG levels against these 4 viruses in a validation cohort of JDM patients recruited in the Netherlands during 2015-2023. Among these 4 viruses tested, we confirmed significantly greater seropositivity to RSV B in JDM patients compared to pediatric controls (p-adj<0.0001) (**Fig.6D**). As vaccination against RSV is only currently available for elderly populations and pregnant women [^43^], these findings suggest that JDM patients had a higher incidence of RSV B infection compared to pediatric controls of the same age.

## DISCUSSION

The unique design of this study, applying in-depth immunological analysis to JDM patients at disease onset, allowed us to implicate the dsRNA sensing pathway as a cause of dysregulated type I IFN responses. Single-cell transcriptomic and chromatin accessibility analyses further revealed functional and epigenetic alterations within the myeloid compartment, including enhanced IFN-I–related programs together with altered antiviral sensing signatures. Furthermore, we provide evidence of RNA viral infection as a potential trigger of disease, consistent with dsRNA sensing pathway dysregulation.

At disease onset, in the absence of prior treatment, JDM patients showed an overactivation of the type I interferon pathway characterized by over production of type I interferon protein subtypes and ISGs. Stimulation of patient blood with dsRNA failed to increase the type I IFN response, while stimulation with a ssRNA agonist (R848) induced production of type I IFN and ISGs similar to age matched controls. Poly(I:C), the dsRNA mimic used in our study can be sensed by multiple sensors. However, of these viral sensors only *IFIH1* (encoding MDA5 protein) showed differential expression compared to controls. No differences in other viral sensors such as TLR3, RLRs or cGAS were observed.

Interestingly among the myositis-specific antibodies (MSAs) that have been proposed to define distinct clinical subgroups of JDM patients [^44,45^], anti-MDA5 antibodies are associated with mild muscle involvements and severe skin manifestations [^46^]. However, the specific contribution of anti-MDA5 antibodies to JDM pathophysiology remains poorly understood. A recent study has shown that myositis aab can be internalized into muscle cells and exert pathogenic effects, including activation of IFN-I signaling by anti-MDA5 aab [^47^]. Among the new onset JDM patients in this study only 3 out of 11 were anti-MDA5 positive, suggesting that the biased response to dsRNA stimulation that we observed is not caused by these autoantibodies. In contrast our single-cell transcriptomic analyses showed that at diagnosis, myeloid cells exhibited transcriptional remodeling preferentially targeting interferon pathway engagement. Enrichment of IRF-binding motifs within differentially accessible chromatin regions in myeloid cells suggests altered accessibility of IRF-associated regulatory elements, potentially uncoupling downstream IFN-I programs despite altered upstream antiviral sensing. This pattern may reflect a pre-activated interferon state in myeloid cells, potentially driven by prior or ongoing immune stimulation, in which sustained IFN-I signaling coexists with impaired responsiveness to specific nucleic acid triggers. Together, these findings support a role for epigenetic remodeling in shaping antiviral sensing and interferon-related programs in JDM myeloid cells. Wilkinson *et al.* previously reported that monocytes are major contributors to interferon-driven pathology in JDM [^48^]. In this study mitochondrial abnormalities in CD14⁺ monocytes were reported to promote the release of oxidized mitochondrial nucleic acid, which induce interferon-stimulated gene expression [^48^].

It has long been speculated that viral infection may be a potential trigger of JDM [^24,41,42^], but providing evidence for this in a rare pediatric disease has been a major challenge. Therefore a major finding of our study is the direct detection of SARS-CoV-2 within muscle tissue at JDM onset. These observations support a model in which localized viral RNA within affected muscle may contribute either directly or indirectly, to the observed tissue damage. Nonetheless the precise significance of the detected viral RNA and whether it reflects transient exposure, residual material or a broader pathological feature requires further investigation. As JDM predates the Covid-19 pandemic, it was important to provide additional evidence of viral infection implication in JDM. For this we took advantage of retrospective biobanked samples which showed JDM patients with significantly higher seropositivity for RSV B in both discovery and validation cohorts. It is interesting to note that among the 27 pathogens tested, only antibodies against RSV B, another RNA virus, were significantly enriched in both cohorts These combined findings, coupled with the defective response of the dsRNA sensing pathway observed at diagnosis, strongly support the implication of RNA viral infection in JDM pathogenesis.

Altered activation of nucleic acid sensors, and viral infectious triggers have also been reported in other complex diseases associated with a type I IFN signature. In systemic lupus erythematosus (SLE), TLR7 was shown to promote inflammation and autoantibody production, while TLR9 played a regulatory role [^49,50^]. More recently, Younis *et al.* provided mechanistic evidence that Epstein–Barr virus (EBV) can infect and reprogram autoreactive B cells in SLE, promoting antigen-presentation programs together with transcriptional and epigenetic remodeling that amplify autoimmune responses [^20^]. These findings support the concept that viral exposure may not only trigger transient interferon responses but also durably reshape immune cell programs involved in disease pathogenesis. In this context, the transcriptional and epigenetic alterations observed in the myeloid compartment in JDM further support the possibility that viral exposure may modulate nucleic acid sensing programs and contribute to the establishment or maintenance of IFN-I–driven inflammation.

A limitation of this study is the small cohort size due to the rare nature of this disease, as proteomic and transcriptomic analyses were based on 11 patients at diagnosis, single cell experiments on one patient and RNAscope on 2. Additionally, prospective inclusion of new onset patients occurred during the Covid-19 pandemic, potentially resulting in recruitment biases. The delay between symptom onset and clinical diagnosis also complicates IgG level interpretation, given variability in IgG persistence across patients and in response to different microbes. This may explain why only the elevated RSV-B seropositivity was confirmed in our replication cohort.

In summary we provide new understanding of dysregulated immunological pathways in JDM, and intriguing evidence in support of a viral trigger. These findings may guide the future development of new therapies targeted to this specific pathway, and provide supportive evidence for preventative strategies such as childhood vaccination against common viral infections.

## MATERIALS AND METHODS

### Study Design

This study aimed to investigate the mechanisms underlying the pathogenesis of juvenile dermatomyositis (JDM). The objectives were to identify mechanisms leading to type I interferon (IFN-I) dysregulation, as well as potential triggers contributing to disease onset. No formal power analysis was performed to determine sample size, which was based on the availability of clinical samples given the rarity of the disease. No predefined stopping rules were applied. All samples meeting inclusion criteria were included in the analysis.

Patients newly diagnosed with JDM were prospectively recruited at their first medical visit between December 2021 and September 2023 from three French pediatric rare disease referral centers (AP-HP Necker–Enfants Malades, AP-HP Robert Debré, and AP-HP Kremlin-Bicêtre). Diagnosis was established according to the European Neuro Muscular Center (ENMC) classification criteria for idiopathic inflammatory myopathies. All samples were collected after informed consent and approved by the Comité de Protection des Personnes (N° EudraCT: 2018-A01358-47). Pediatric healthy controls, including those from the ATRACTion cohort for serological analyses, were recruited from AP-HP Necker–Enfants Malades and had no history of immune-related disease. Where applicable, samples were age-matched across cohorts to ensure comparability. Samples with missing data or technical issues were excluded prior to analysis. No additional outlier exclusion criteria were applied.

Primary endpoints were defined based on established biological hypotheses, particularly focusing on IFN-I–related immune signatures. An initial exploratory phase characterized baseline cytokine and transcriptomic profiles, which informed subsequent hypothesis-driven analyses. Statistical comparisons were performed using non-parametric tests, and p-values were adjusted for multiple testing using the FDR method. Each measurement represents an independent biological sample. No technical replicates were performed unless otherwise specified. This was a prospective observational cohort study. No randomization or blinding was performed. Samples were collected and processed under standardized conditions and stored after initial processing for batched analyses. We performed different experimental protocols (detailed below) tailored to the requirement of each analyses. Data analysis was conducted using pre-defined statistical approaches.

### Patient clinical characteristics

In this study we included 19 patients at disease diagnosis. Each patient was tested for myositis-specific antibodies (MSAs). MSAs and experimental information are described in **Table S3**.

### TruCulture whole blood *ex vivo* stimulation

TruCulture tubes (Rules Based Medicine - Q^2^ Solutions Company, Austin, USA) with the following selected agonists were used; Poly(I:C) (20μg/ml) to mimic double-stranded RNA viruses activating the TLR3/IFIH1 pathway and R848 (1μM) to mimic single-stranded RNA viruses activating TLR7 and TLR8. For each patient and healthy control, an unstimulated tube (Null) served as a control. As previously described, 1ml of whole blood collected in Lithium Heparin tubes was added to each TruCulture tube within 1 hour of blood collection [^52^] and incubated at 37°C in a dry block incubator for 22 hours. Following incubation, TruCulture supernatants were aliquoted and stored at −80°C for subsequent cytokine analyses. 2ml of TRIzol LS Reagent (Life Technologies) was added to the cell pellet for later RNA extraction and transcriptomic analysis as previously described [^53^].

### Cytokine assays

TruCulture supernatants were assessed for secreted cytokines using the Human XL 46-plex panel (R&D Systems) via bead-based ELISA (Luminex). All concentrations were measured in pg/ml – except CXCL2, CCL5, PDGF-AA, PDGF-AB/BB in ng/ml. IFN-α subtypes and IFN-β were measured in a duplex digital ELISA Simoa (Quanterix) as previously described [^25,54^].

### RNA extraction

Total RNA was extracted from TRIzol lysate of TruCulture (detailed below cell pellets) using a modified protocol of the NucleoSpin RNA kit (Macherey-Nagel, Dueren, Germany) as previously described [^53^]. Briefly, 600μl of TRIzol lysate were resuspended with 900μl of absolute ethanol and transferred to NucleoSpin columns. After 30 seconds of 11,000g centrifugation, the columns were washed with MW1 (1 time) and MW2 (2 time). RNA was then eluted in 60μl of RNAse Free Water. RNA concentration was measured using the Qubit RNA HS Assay Kit (Life Technologies) according to the manufacturer’s protocol. RNA integrity ratio (260/230 value) was performed using the Nanodrop spectrophotometer (ThermoFischer Scientific). After quality controls, samples were stored at -80°C.

### Gene expression analysis

Bulk RNA barcoding and sequencing (BRB-seq) protocol was used for the transcriptomic study of all samples, as previously described [^55^] by Alithea Genomics, Epalinges, Switzerland. Unique molecular identifier (UMIs) counts were exported in csv files and imported into R (4.1.2). Raw BRB-seq data were processed directly in DESeq2 package for differential expression analysis. For each comparison (e.g., JDM stims (Poly:IC/R848) vs. JDM Null, Controls stims (Poly:IC/R848) vs. Controls Null), an independent filtering implemented in the DESeq2 package was applied to automatically remove genes with low mean normalized counts that lacked sufficient power for reliable statistical testing. For Poly(I:C), 14,797 genes were retained in controls and 10,463 in JDM patients, for R848, 16,035 genes were retained in controls and 14,178 in JDM patients. Principal component analysis (PCA), a multidimensional scaling method, was performed using prcomp method from stats packages, and was plotted using plotly and ggplot2 libraries. Differentially expressed genes were explored using DESeq2 package with a threshold at <0.05 for the adjusted p-value (calculated from Wald test and corrected using Benjamini and Hochberg method) and <|1| for the log2FoldChange. Pathway enrichment analysis was assessed with clusterProfiler package in R using Gene Ontology (GO) database as gene sets. We performed weighted gene correlation network for analysis (WGCNA) using the ‘wgcna’ package in R. (4.1.2). Once counts were normalized, we picked a soft threshold power of 9 to create the network and identify gene modules. We next calculated the Eigangene for each module to determine if specific modules were more correlated with specific conditions.

### Mass Cytometry

Mass cytometry (CyTOF) was carried out to characterize circulating immune cells in 6 JDM patients at diagnosis and 5 paediatric controls (**Table S11**). From 300μl of whole blood, we labelled several membrane markers with metal-isotope-tagged antibodies for different immune cells populations using two different panels (**Table S12**). Cells were fixed with Proteomic Stabilizer PROT1 (Smart Tube Inc., Las Vegas, USA) and stored at -80°C for later batch analysis. Once samples were batched and thawed, red blood cells were lysed using Thaw Lyse 1000X (Smart Tube Inc., Las Vegas, USA). After resuspension in PBS supplemented with 2% formaldehyde (Thermo Scientific) and 125 nM of Cell-ID™ Intercalator-Ir (Standard BioTools), samples were incubated overnight at 4°C and then frozen at -80°C until analysis with a Helios mass cytometer (Standard BioTools). Datasets were analysed using OMIQ software (Dotmatics, Boston, USA). Analysis was performed on CD45+ and CD66b- viable cells only. All samples followed the same gating strategy to identify immune cell populations and subpopulations (**Fig.S1**).

### Single-cell RNA sequencing (scRNA-seq) and single-nuclei ATAC sequencing (snATAC-seq)

Peripheral blood mononuclear cells (PBMCs) were isolated from one JDM patient at disease onset and two age- and sex-matched healthy controls, and cryopreserved at -150°C. The day prior to the experiments, cells were thawed and left to rest overnight (∼ 18 hours) at 37°C in RPMI medium. Cell viability after resting was > 70%. For each of the samples, a Chromium GEM-X v4 Universal 3ʹ Gene Expression (User Guide CG000731_RevB) library (GEX) and a Chromium Next GEM v2 Single Cell ATAC (User Guide CG000496_RevC) library (ATAC) (10x Genomics) were generated in parallel, targeting 20,000 cells or nuclei per library. Nuclei were isolated from an aliquot of 300,000 cells from each of the samples, following 10x Genomics guidelines (Demonstrated Protocol CG000169_Rev E, 4-min lysis time). GEX and ATAC libraries quality and fragment size distribution were assessed using a Bioanalyzer 2100 (Agilent), and concentrations were determined using a Qubit 2.0 fluorometer (Thermo Fisher Scientific). All processed libraries were sequenced on a NextSeq 2000 (Illumina) using P4 (100 cycles, 1.8B reads) flow cells and paired-end sequencing parameters recommended by 10x Genomics. GEX and ATAC raw sequencing data were processed using *CellRanger* (v9.0.0) and *CellRanger ATAC* (v2.1.0) (10x Genomics), respectively. Reads were aligned to the human reference genome (GRCh38) and feature-barcode matrices were generated using default parameters for each library.

Pre-processing of the GEX modality libraries was performed using *scanpy* (v1.11.85) [^56^]. In each library, cell barcodes with over 20% mitochondrial reads or less than 1,000 detected genes were filtered out. Genes not detected in at least 10 cells were also excluded. Doublets were detected and removed using the *scrublet* [^57^] implementation built into *scanpy* with default parameters. All three libraries were then concatenated into a single *AnnData* (0.10.9) object. UMI counts were log-normalized, and the 2,000 most variable genes were considered to compute the 50 first principal components of the data, compute a 25-nearest-neighbor graph, cluster barcodes using *scanpy*’s implementation of the Leiden algorithm and compute UMAP coordinates. Cell-type identities were inferred using *celltypist* (v1.7.1) [^58^] with an in-house set of transcriptional markers for 42 PBMC types. Given these cell-type identities, barcode sets were aggregated into myeloid cells, as well as lymphoid B, CD4⁺ T, CD8⁺ T and NK cells.

The ATAC modality libraries were pre-processed using *SnapATAC2* (v2.7.1) [^59^]. In each library, nucleus barcodes with a fragment count lower than 1,000 or higher than 100,000, or a TSS enrichment score below 10 were filtered out. We then computed paired-insertion counts across 500-base-pair-wide genomic bins for each barcode, excluding a set of genomic regions blacklisted by ENCODE (ENCFF356LFX) [^60^]. Doublet barcodes were detected and filtered out using the *scrublet* implementation built into *SnapATAC2* with default parameters.

For each nucleus barcode and each gene considered in the GEX modality, we also computed a ‘gene activity score’ as a function of chromatin accessibility along the gene body, extending up to 2,000 base-pairs upstream of the TSS. Based on these gene activity scores, we used the *scVI* (v0.14.6) [^61^] variational auto-encoder to learn a 50-dimensional latent representation of the GEX and ATAC data points in each sample. To this end, we trained a neural network with 5 hidden layers and 256 nodes per layer; training stopped after 50 validation epochs with no improvement in evidence lower bound (ELBO). We then used *scANVI* (v0.14.6) [^62^] to transfer immune-lineage labels from the GEX to the ATAC data points from each sample. Following label-transfer, we kept only confidently labeled data points (posterior probability > 0.8) and removed clusters of ATAC data points bearing a label not concordant with neighboring data points in UMAP coordinates (e.g., nuclei-barcode clusters from one sample annotated as myeloid but projecting with CD4⁺ T nuclei from other samples).

We then concatenated all three ATAC libraries into a single *AnnData* (0.10.9) object and used the *MACS3* implementation of *SnapATAC2* to call a set of 202,770 accessibility ‘peaks’ or open-chromatin-regions common to all samples across all immune lineages. We assigned each peak to a type of cis-regulatory element based on the ChromHMM ‘full-stack’ annotation from [^63^]. To infer regulatory activity-by-contact, we overlapped our peaks with sets of regions in predicted contact with gene promoters in each immune cell type [^39,40^].

Differential gene expression and chromatin accessibility tests between the JDM onset patient and the age-matched controls were performed with *Scanpy* and *SnapATAC2* respectively, considering cells or nuclei assigned to each immune lineage. Differentially expressed genes show an absolute log-fold change (logFC) > 0.5 at a false discovery rate (FDR) < 0.01; differentially accessible peaks show an absolute log_2_FC > 0.5 at an FDR < 0.01. Functional enrichment tests in the GEX modality were performed using *GOSeq* (v1.44.0) ^64^ on differentially expressed genes, stratified by effect direction (i.e. upregulated in the JDM patient or the controls). Functional enrichment tests in the ATAC modality were performed using *GOSeq* on the sets of genes linked through activity-by-contact to differentially accessible peaks, stratified by effect direction. Transcription factor motif enrichment analyses were performed using *SnapATAC2*, considering differentially accessible regions against all accessible regions in each lineage, using a list of transcription factor motifs curated by the CIS-BP database [^65^].

### Antibody serology test

A bead-based serological assay (Luminex) was used to detect immunoglobulin G (IgG) levels in the first plasma of each patient stored in our biobank and paediatric controls, as previously described [^66,67^]. For antibody serology analyses, samples were selected from our retrospective serum biobank comprising patients recruited in Parisian pediatric rare disease referral centers (initial cohort) and from a national JDM study including six University Medical Centers in the Netherlands (validation cohort). All samples were collected following informed consent procedures (as described above for the French cohort) and with approval from the institutional ethics committee of the University Medical Center Utrecht, the Netherlands (NL47875.041.14). Serum samples were stored at −80 °C until IgG analysis. For each patient, the earliest available sample closest to disease onset was selected. The French cohort included 68 patients with samples collected between 2017 and 2023, while the Dutch validation cohort comprised 62 patients with samples collected between 2015 and 2023. Antibodies were measured simultaneously against 27 different microbial pathogens (**Table S9**). For each antigen tested, we calculated a related antibody unit (RAU), as well as a serostatus cutoff as previously defined and described in a larger data set [^67^].

### RNAscope in situ hybridization on muscle biopsies

RNAscope in situ hybridization (ISH) was performed on skeletal muscle biopsies obtained from newly diagnosed JDM patients during the Covid-19 pandemic (n = 2; July 2020 and March 2021) and from one patient with JDM at disease onset prior to the Covid-19 pandemic (n = 1). Ten µm sections of fresh-frozen samples were mounted on positively charged SuperFrost Plus slides (Fisher Scientific) and fixed with 4% paraformaldehyde. Muscle biopsies were embedded in tragacanth gum, mounted on cork to maintain orientation, and snap-frozen in isopentane cooled in liquid nitrogen, rather than embedded in OCT. RNAscope® ISH was used to localize SARS-CoV-2-N in human muscle. Probes were developed and manufactured by Bio-Techne (Minneapolis, Minnesota, USA). Details of the optimized protocol are available online on the ACDbio website. The samples were subjected to a protease-based pretreatment. Slides were then incubated with RNAscope® probes including a negative control probe (#320871), a positive control probe (Hs-PPIB, #313901), and V-SARS-CoV-2-N-O6-C1 probe (#1862171-C1). After three amplification steps, a fluorophore was applied: TSA Vivid 650 (#323273) for C1. Slides were counterstained with DAPI and imaged using a ZEISS AxioScan 7 digital slide scanner with ZEN software (version 3.9.101) (Zeiss®, Oberkochen, Germany).

### Statistical analysis

Significant corrected p-values are indicated by star symbols as follows: *=p < 0.05; **=p < 0.01; ***=p < 0.001; ****=p<0.0001. Cytokine levels were log2 normalized to account for the high variability and wide range in the original cytokine measurements. Mann-Whitney tests were employed to compare cytokine levels between unstimulated JDM at diagnosis versus unstimulated controls (two-way tests), p-values were corrected using Benjamini-Hochberg method with p-adj significance threshold at <0.05. Correlations between gene expression levels were performed using the Sperman test, with p-values corrected (p-adj) using the False Discovery Rate (FDR). Heatmap was generated using pheatmap package. A Kruskal-Wallis test followed by a post-hoc Dunn’s multiple comparisons test was used to assess significant differences between JDM patients and controls in response to TLR stimulations (multiple group tests). Dotplots were generated with GraphPad Prism (10.1.1). Contingency tests with FDR correction were used to compare serostatus between control and patient groups. All plots were generated using either R (4.1.2), GraphPad Prism (10.1.1).

## ACKNOWLEDGEMENTS

Fundings and declarations

This study was supported by funding from the ANR (Agence Nationale de Recherche), grant number ANR-21-CE17-0025 JDMINF2, ANR-25-CE14-3602 SUNSHINE and from DMU MEFADO Département Médico-Universitaire Médecine de l’enfant et de l’adolescent Necker Hospital Enfants-Malades, AP-HP, Paris, France. DD acknowledges support from the Institut Pasteur Explore programme. The Attraction cohort was supported by an RHU grant (ref ANR-18-RHUS-0010). The authors would like to thank the Comité des fêtes of Kervardel and its generous donators for their financial supports. We also would like to thank Sabrina Guermah for her technical assistance. Graphical abstract was created using Biorender.com.

## Authorship contributions: CRediT

**Data acquisition: TRJM, YA, YYZ, VB, CAV, FDon., FDub., BP, FR, YT, SRV, CBru., BH, AV, DHin., TT, CMC**

**Clinical recruitments: SRV, CBod., MLF, BF, PQ, FJA, ES, AS, AVRK, AWM, FVW, FRL, MJ, IM, BBM, CG**

**Resources: SRV, FJA, FRL, MJ, MH, DHar., MW, LQM, MPR, DD**

**Project conceptualization, administration and supervision: DD, MPR, CG, BBM**

**Data analysis and interpretation: TRJM, YA, VB, FDub., EV, AB, DHar., MW, MPR, DD**

**Writing-original draft: TRJM, YA, DD, MPR**

**Writing-review and editing: TRJM, YA, YYZ, VB, CAV, FDon., FDub., BP, FR, YT, SRV, EV, AB, CBod., CBru., MLF, BF, BH, PQ, FJA, ES, AS, AV, AVRK, AWM, FVW, FRL, MJ, DHin., TT, CMC, MH, DHar., MW, LQM, IM, BBM, CG, MPR, DD**

## Data availability

The datasets generated during and/or analysed during the current study are available from the corresponding author on reasonable request.

## Ethics approval

All samples were collected after informed consent and approved by the Comité de Protection des Personnes (N° EudraCT: 2018-A01358-47 in France and NL47875.041.14 in the Netherlands).

## LIST OF SUPPLEMENTARY MATERIALS

**Tables S1 to S12 for multiple supplementary tables listed below:**

**Table S1.** Enriched pathways upregulated from Gene Ontology (GO) database at baseline in JDM at diagnosis compared to pediatric controls. ***See Excel File TAB_S1 GO JDM_NULL HD_NULL.xlsx***

**Table S2.** Enriched pathways downregulated from Gene Ontology (GO) database at baseline in JDM at diagnosis compared to pediatric controls.

**Table S3.** Myositis-specific antibodies and experiments table of JDM at diagnosis cohort.

**Table S4.** Adjusted p-values results of Kruskal-Wallis test and post-hoc Dunn’s test of 46 cytokines measured after TLRs stimulation. Not applicable indicates cytokines that were below the limit of detection.

**Table S5.** Differentially expressed genes across immune cell lineages in the JDM patient. ***See Excel File TAB_S5 DE P2_HD.xlsx***

**Table S6.** GO enrichment analysis of differentially expressed genes between JDM patient and healthy donors. ***See Excel File TAB_S6 DE P2_HD GOSEQ.xlsx***

**Table S7.** Custom gene sets for single-cell analysis with Gene_ID content.

**Table S8.** Transcription factor motif enrichment in differentially accessible chromatin regions (JDM vs controls). ***See Excel Files TAB_S8A DA JDM_HD ; TAB_S8B DA JDM_HD TFBM_ENRICH.xlsx***

**Table S9.** Microbial Pathogen-Associated Antigens Detected in IgG Assay.

**Table S10.** Adjusted p-values from contingency testing of IgG plasmatic levels in the initial JDM cohort.

Table S11. Immune Cell Population Phenotypic Markers Used in Mass Cytometry (CyTOF) Analysis.

**Table S12.** Fluorochrome-conjugated antibodies for membrane antigen staining in Mass Cytometry (CyTOF) analysis.

**Figures S1 to S7 for multiple supplementary figures listed below:**

**Figure S1. Correlations between ISGs gene counts and *IFIH1*.** Correlation plot between *IFIH1* gene count and well-described interferon stimulated genes (ISGs) count. Each dot represents a patient. n=11 JDM at diagnosis. p-values are corrected (p-adj) using False Discovery Rate (FDR) method.

**Figure S2. Number of Differentially Expressed Genes (DEGs) upregulated and downregulated according to stimulation and patients or controls.** Venn diagram representing the number of DEGs after Poly(I:C) (green) or R848 (red) compared to unstimulated condition in patients (JDM) and controls.

**Figure S3. Enriched pathways upregulated in controls and JDM at diagnosis after R848 stimulations compared to their respective baseline profiles.** Pathways are a list of genes from GO enrichment database. The count represents the number of genes from DEGs that are found in a specific Gene Ontology (GO) term. The GeneRatio is the proportion of our DEGs associated with a particular GO term relative to the total number of our DEGs.

**Figure S4. Dendrogram from Weighted correlation network analysis (WGCNA) reveals distinct genes modules.** This dendrogram shows the hierarchical clustering of genes based on topological overlap in gene expression data, generated using the WGCNA package. Each branch represents a module of color (group of genes with similar expression patterns across the samples). On the y-axis, the height indicates the divergence in gene expression between clusters, with lower values indicating greater similarity.

**Figure S5. Enriched pathways analysis from the “turquoise”, “greenyellow” and “red gene modules of the WGCNA analysis.** Pathways are a list of genes from GO enrichment database. The count represents the number of genes from DEGs that are found in a specific Gene Ontology (GO) term. The GeneRatio is the proportion of our DEGs associated with a particular GO term relative to the total number of our DEGs.

**Figure S6. Number of JDM at diagnosis recruited during the study compared at the number expected before the start of the study based on country incidence.**

**Figure S7. Histogram showing the distribution of seropositive (POS) and seronegative (NEG) individuals among controls (grey) and JDM patients (purple) for SARS-CoV-2 Wuhan RBD.**

