## supplemental tables and figures for "Dysregulated dsRNA sensor signaling and viral infection during onset of pediatric autoimmune interferonopathy"

### SUPPLEMENTARY TABLES AND FIGURES

**Table S1. Enriched pathways upregulated from Gene Ontology (GO) database at baseline in JDM at diagnosis compared to pediatric controls.**

***See Excel File TAB_S1__GO__JDM_NULL__HD_NULL.xlsx***

| Description | p.adjust (BH) | geneID |
|---|---|---|
| gamma-delta T cell activation | 0,01741 | ITK/LEF1/CD3G/CCR9/TRDC |
| ribosome biogenesis | 0,01939 | TFB1M/RPL6/C1QBP/AIRIM/RPF1/UTP25/RPL5/RPS6/URB1/RPS8/LYAR/MTERF3/RIOX2/RPS23 |
| T cell receptor signaling pathway | 0,02347 | ITK/**CD8A**/CD3G/CD8B/THEMIS/CD28/SH2D1A/TRDC/LAT |
| T cell differentiation | 0,02347 | RORA/CRTAM/ITK/BCL11B/RPS6/LEF1/RORC/CAMK4/**CD8A**/CD3G/THEMIS/CCR9/CD28 |
| mononuclear cell differentiation | 0,04433 | RORA/ITM2A/CRTAM/ITK/GPR68/BCL11B/MYC/RPS6/LEF1/RORC/CAMK4/**CD8A**/CD3G/THEMIS/CCR9/CD28 |

**Table S2. Enriched pathways downregulated from Gene Ontology (GO) database at baseline in JDM at diagnosis compared to pediatric controls.**

| Patients | Exepriments | MSAs |
|---|---|---|
| P1 | TruCulture, IgG dosage | MSA Neg |
| P2 | TruCulture, IgG dosage | TIF1γ+ |
| P3 | TruCulture, IgG dosage | Mi-2+ |
| P4 | TruCulture, IgG dosage | MSA Neg |
| P5 | TruCulture, IgG dosage | Mi-2+ |
| P6 | TruCulture, IgG dosage | NXP2+ |
| P7 | TruCulture, IgG dosage | MDA5+ |
| P8 | TruCulture, IgG dosage | NXP2+ |
| P9 | TruCulture, IgG dosage | TIF1γ+ |
| P10 | TruCulture, IgG dosage | MDA5+ |
| P11 | Mass cytometry, TruCulture, IgG dosage | MDA5+ |
| P12 | Mass cytometry | TIF1γ+ |
| P13 | Mass cytometry, IgG dosage | NXP2+ |
| P14 | Mass cytometry, IgG dosage | MSA Neg |
| P15 | Mass cytometry | Mi-2+ |
| P16 | Mass cytometry | MSA Neg |
| P17 | Single cellular experiments | MDA5+ |
| P18 | RNAScope | MDA5+ |
| P19 | RNAScope | TIF1γ+ |
| **MSAs: Myositis-Specifc Antibodies, TIF1γ: transcription intermediary factor 1-γ, MDA5: melanoma differentiation-associated protein 5, NXP2: nuclear matrix protein 2** | | |

**Table S3. Myositis-specific antibodies and experiments table of JDM at diagnosis cohort.**

| Cytokine | Kruskal Wallis Test Controls | Kruskal Wallis Test Patients | Adjusted p-values Controls Null vs Poly(I:C) | Adjusted p-values JDM at diagnosis Null vs Poly(I:C) | Adjusted p-values Controls Null vs R848 | Adjusted p-values JDM at diagnosis Null vs R848 |
|---|---|---|---|---|---|---|
| **TNFα** | 0,00001 | 0,00002 | 0,01885 | 0,46608 | 0,00000 | 0,00010 |
| **TNFβ** | 0,00009 | 0,00001 | 0,68127 | 0,13771 | 0,00039 | 0,00100 |
| **TRAIL** | 0,01745 | 0,22050 | 0,42500 | Not Applicable | 0,01313 | Not Applicable |
| **IFNα** | 0,00000 | 0,00009 | 0,00821 | 0,22751 | 0,00000 | 0,00007 |
| **IFNβ** | 0,00000 | 0,00000 | 0,00056 | 0,03567 | 0,00000 | 0,00001 |
| **IFNγ** | 0,00000 | 0,00296 | 0,00274 | 0,44557 | 0,00003 | 0,00621 |
| **IL-1α** | 0,00000 | 0,00002 | 0,00325 | 0,27899 | 0,00000 | 0,00002 |
| **IL-1β** | 0,00000 | 0,00000 | 0,08150 | 0,28062 | 0,00000 | 0,00000 |
| **IL-1ra** | 0,00001 | 0,00001 | 0,01150 | 0,21908 | 0,00000 | 0,00001 |
| **IL-2** | 0,00028 | 0,00000 | 0,11457 | 0,03334 | 0,00590 | 0,00136 |
| **IL-3** | 0,74511 | 0,49655 | Not Applicable | Not Applicable | Not Applicable | Not Applicable |
| **IL-4** | 0,00056 | 0,25793 | 0,33792 | Not Applicable | 0,01598 | Not Applicable |
| **IL-5** | 0,06910 | 0,03997 | Not Applicable | 0,19534 | Not Applicable | 0,01371 |
| **IL-6** | 0,00000 | 0,00002 | 0,01429 | 0,28502 | 0,00000 | 0,00002 |
| **IL-7** | 0,12336 | 0,09504 | Not Applicable | Not Applicable | Not Applicable | Not Applicable |
| **IL-8** | 0,00006 | 0,00001 | 0,18241 | 0,24041 | 0,00007 | 0,00001 |
| **IL-9** | 0,01545 | 0,11230 | 0,08862 | Not Applicable | 0,21190 | Not Applicable |
| **IL-10** | 0,00000 | 0,00002 | 0,01964 | 0,48643 | 0,00000 | 0,00019 |
| **IL-12p70** | 0,00028 | 0,02918 | 0,01034 | 0,09739 | 0,00009 | 0,00813 |
| **IL-13** | 0,00021 | 0,00003 | 0,43363 | 0,07261 | 0,00308 | 0,01409 |
| **IL-15** | 0,03902 | 0,58081 | 0,36241 | Not Applicable | 0,05491 | Not Applicable |
| **IL-17A** | 0,00236 | 0,31752 | 0,13014 | Not Applicable | 0,07390 | Not Applicable |
| **IL-17E** | 0,00229 | 0,04423 | 0,42017 | 0,37294 | 0,14398 | 0,20625 |
| **IL-33** | 0,00015 | 0,03264 | 0,05872 | 0,13660 | 0,00003 | 0,00983 |
| **CXCL10** | 0,00019 | 0,00012 | 0,00256 | 0,23189 | 0,00007 | 0,00019 |
| **CCL2** | 0,00000 | 0,00001 | 0,01228 | 0,30467 | 0,00000 | 0,00001 |
| **CCL3 MIP-1α** | 0,00002 | 0,00002 | 0,07479 | 0,58432 | 0,00002 | 0,00013 |
| **CCL4 MIP-1β** | 0,00000 | 0,00001 | 0,00420 | 0,22216 | 0,00000 | 0,00000 |
| **CCL20 MIP-3α** | 0,00001 | 0,00000 | 0,21452 | 0,28158 | 0,00059 | 0,00018 |
| **MIP-3β** | 0,00001 | 0,01968 | 0,28972 | 0,10882 | 0,00038 | 0,13602 |
| **CXCL1 GRO-α** | 0,00001 | 0,00008 | s | 0,32785 | 0,00837 | 0,00144 |
| **CXCL2 GRO-β** | 0,00090 | 0,00089 | 0,50019 | 0,47152 | 0,00267 | 0,00127 |
| **CCL5 RANTES** | 0,05546 | 0,24562 | Not Applicable | Not Applicable | Not Applicable | Not Applicable |
| **FLT-3-ligand** | 0,02216 | 0,11140 | 0,10509 | Not Applicable | 0,00608 | Not Applicable |
| **G-CSF** | 0,00037 | 0,02730 | 0,43778 | 0,16894 | 0,00180 | 0,16750 |
| **GM-CSF** | 0,01606 | 0,40720 | 0,40610 | Not Applicable | 0,05457 | Not Applicable |
| **EGF** | 0,01407 | 0,18008 | 0,20309 | Not Applicable | 0,12084 | Not Applicable |
| **FGF-basic** | 0,00021 | 0,00056 | 0,14652 | 0,05326 | 0,03185 | 0,04670 |
| **PDGF-AA** | 0,13487 | 0,15428 | Not Applicable | Not Applicable | Not Applicable | Not Applicable |
| **PDGF-AB/BB** | 0,02977 | 0,04351 | 0,52653 | 0,25118 | 0,46190 | 0,12022 |
| **VEGF** | 0,04022 | 0,15651 | 0,12937 | Not Applicable | 0,28472 | Not Applicable |
| **PD-L1/B7-H1** | 0,00164 | 0,13168 | 0,25284 | Not Applicable | 0,00262 | Not Applicable |
| **CD40 Ligand** | 0,00197 | 0,09806 | 0,15573 | Not Applicable | 0,08116 | Not Applicable |
| **Eotaxin** | 0,13890 | 0,34375 | Not Applicable | Not Applicable | Not Applicable | Not Applicable |
| **Granzyme-B** | 0,00001 | 0,06518 | 0,00584 | Not Applicable | 0,00010 | Not Applicable |
| **TGFα** | 0,07932 | 0,09457 | Not Applicable | Not Applicable | Not Applicable | Not Applicable |

**Table S4. Adjusted p-values results of Kruskal-Wallis test and post-hoc Dunn's test of 46 cytokines measured after TLRs stimulation. Not applicable indicates cytokines that were below the limit of detection.**

**Table S5. Differentially expressed genes across immune cell lineages in the JDM patient.**
***See Excel File TAB_S5__DE__P2_HD.xlsx***

**Table S6. GO enrichment analysis of differentially expressed genes between JDM patient and healthy donors.**
***See Excel File TAB_S6__DE__P2_HD__GOSEQ.xlsx***

| Geneset Name | Gene_ID |
|---|---|
| Crow IFN-I Signature | IFI27, IFI44L, IFIT1, RSAD2, ISG15, SIGLEC1 |
| Pathogen Sensors | TLR1, TLR2, TLR3, TLR4, TLR5, TLR6, TLR7, TLR8, TLR9, NOD1, NOD2, NAIP, NLRP1, AIM2, MB21D1, IFI16, DDX41, DDX58, IFIH1, ZBP1, OAS1, OAS2, OAS3 |
| Viral Sensors | AIM2, CGAS, IFI16, DDX41, TLR3, TLR7, TLR8, RIGI, IFIH1, ZBP1, OAS1, OAS2, OAS3 |
| RNA Viral Sensors | TLR3, TLR7, TLR8, RIGI, IFIH1, ZBP1, OAS1, OAS2, OAS3 |

**Table S7. Custom gene sets for single-cell analysis with Gene_ID content.**

**Table S8. Transcription factor motif enrichment in differentially accessible chromatin regions (JDM vs controls).**
***See Excel Files TAB_S8A__DA__JDM_HD.xlsx ; TAB_S8B __DA__JDM_HD__TFBM_ENRICH.xlsx***

| Microbial Pathogens | Antigens |
|---|---|
| Adenovirus | Adenovirus T3 |
| | Adenovirus T5 |
| | ADE5 |
| | ADE40 |
| Bordetella | Bordetella p. Toxin |
| | Bordetella p. FHA |
| Cytomegalovirus | CMV |
| Diphteria | Diptheria Toxin |
| Epstein-Barr Virus | EBV gp125 |
| Echovirus | Echovirus |
| Enterovirus | Enterovirus CoxB3 VP1 |
| Hepatitis A | Hepatitis A |
| Hepatitis E | HEV ORF2 |
| Human Coronavirus OC43 | OC43 NP |
| | OC43 S |
| Human Coronvavirus HK | HKU1 NP |
| | HKU1 S |
| Human Coronvavirus 229E | 229E NP |
| | 229E S |
| Human Coronvavirus NL63 | NL63 NP |
| | NL63 S |
| Human Coronavirus SARS-CoV-2 | SARS-CoV-2 Spike Wuhan |
| | SARS-CoV-2 RBD Wuhan |
| | SARS-CoV-2 Spike Omicron |
| | SARS-CoV-2 RBD Omicron |
| | SARS-CoV-2 NP |
| | SARS-CoV-2 Spike S2 |
| | ME fusion |
| Human papillomavirus | HPV 16 |
| | HPV 18 |
| Influenza A | FluA |
| Measles | Measles lysate |
| | Measles NP |
| Mumps | Mumps lysate |
| | Mumps NP |
| | Mumps |
| Nororvirus | Norovirus GII.4 VLP |
| | Norovirus GII.4 VP1 |
| | Norovirus GII.6 VLP |
| Leptospirosis | Ag Lepto CNR |
| Respiratory Syncytial Virus | Respiratory Syncytial A |
| | Respiratory Syncytial B |
| | RSV gG |
| Rhinovirus | Rhinovirus T1A lysate |
| Rotavirus | Rotavirus VP7 |
| Rubella | Rub VLP |
| Tetanus | Tetanus Toxin |
| | Tet Tox |
| Toxoplasmosis | Toxoplasma gondii SAG2 |
| Varicella | Varicella-Zoster Virus |

**Table S9. Microbial Pathogen-Associated Antigens Detected in IgG Assay.**

| Antigens | Adjusted p-value (FDR) |
|---|---|
| Adenovirus_T3_Dilution | 0,47628 |
| Adenovirus_T5_Dilution | 0,05044 |
| Bordetella_p._Toxin_Dilution | 0,62977 |
| Bordetella_p._FHA_Dilution | 1,00000 |
| CMV_Dilution | 0,30775 |
| Diptheria_Toxin_Dilution | 1,00000 |
| EBV_gp125_Dilution | 0,21774 |
| Echovirus_Dilution | 1,00000 |
| Enterovirus_CoxB3_VP1_Dilution | 1,00000 |
| HEV_ORF2_Dilution | 0,04522 |
| Measles_lysate_Dilution | 0,17511 |
| Mumps_lysate_Dilution | 1,00000 |
| Norovirus_GII.4_VLP_Dilution | 0,62977 |
| Norovirus_GII.4_VP1_Dilution | 0,52322 |
| Norovirus_GII.6_VLP_Dilution | 0,30775 |
| Respiratory_Syncytial_A_Dilution | 1,00000 |
| Respiratory_Syncytial_B_Dilution | 0,01301 |
| RSV_gG_Dilution | 1,00000 |
| Rhinovirus_T1A_lysate_Dilution | 1,00000 |
| Rotavirus_VP7_Dilution | 1,00000 |
| Tetanus_Toxin_Dilution | 0,87911 |
| Varicella_Zoster_Virus_Dilution | 0,62977 |
| Toxoplasma_gondii_SAG2_Dilution | 0,05044 |
| Hepatitis_A_Dilution | 0,27235 |
| Mumps_Dilution | 0,62977 |
| HPV_16_Dilution | 1,00000 |
| HPV_18_Dilution | 0,13598 |
| OC43_NP_Dilution | 1,00000 |
| OC43_S_Dilution | 0,05044 |
| HKU1_NP_Dilution | 0,62977 |
| HKU1_S_Dilution | 0,64014 |
| 229E_NP_Dilution | 0,47628 |
| 229E_S_Dilution | 1,00000 |
| NL63_NP_Dilution | 0,12540 |
| NL63_S_Dilution | 0,43891 |
| FluA_Dilution | 0,02172 |
| ADE5_Dilution | 0,04522 |
| ADE40_Dilution | 0,13598 |
| mumps_NP_Dilution | NA |
| measles_NP_Dilution | 0,05044 |
| Rub_VLP_Dilution | NA |
| Tet_Tox_Dilution | 1,00000 |

**Table S10. Adjusted p-values from contingency testing of IgG plasmatic levels in the initial JDM cohort.**

| Cell Populations | Cell Markers |
|---|---|
| Naive CD4 T cells | CD3+ CD4+ CCR7+ CD45RA+ |
| Central Memory CD4 T cells | CD3+ CD4+ CCR7+ CD45RA- |
| Effector Memory CD4 T cells | CD3+ CD4+ CCR7- CD45RA- |
| Effector CD4 T cells | CD3+ CD4+ CCR7- CD45RA+ |
| Naive CD8 T cells | CD3+ CD8+ CCR7+ CD45RA+ |
| Central Memory CD8 T cells | CD3+ CD8+ CCR7+ CD45RA- |
| Effector Memory CD8 T cells | CD3+ CD8+ CCR7- CD45RA- |
| Effector CD8 T cells | CD3+ CD8+ CCR7- CD45RA+ |
| NK cells | CD14-CD3-CD56+ |
| Transitional B cells | CD3-CD19+CD38hi |
| Naive B cells | CD3-CD19+CD27- |
| Memory B cells | CD3-CD19+CD27+ |
| Monocytes | CD3-CD19-CD14+ |
| Plasmacytoid dendritic cells (pDCs) | CD3-CD19-CD14-HLA-DR+CD123+ |
| Basophils | CD3-CD19-CD14-HLA-DR-CD123+ |

**Table S11. Immune Cell Population Phenotypic Markers Used in Mass Cytometry (CyTOF) Analysis.**

| Antibodies | Label | Clone | Manufacturer |
|---|---|---|---|
| CD45 | 089Y | HI30 | Fluidigm |
| CD196 (CCR6) | 141Pr | G034E3 | Fluidigm |
| CD19 | 142Nd | HIB19 | Fluidigm |
| CD123 | 143Nd | 6H6 | Fluidigm |
| CD38 | 144Nd/172Tm* | 5HIT2 | Fluidigm |
| CD4 | 145Nd | RPA-T4 | Fluidigm |
| CD8a | 146Nd | RPA-T8 | Fluidigm |
| CD66b | 152Sm/169Tm* | 80H3 | Fluidigm |
| CD3 | 154Sm | UCHT1 | Fluidigm |
| CD27 | 155Gd | L128 | Fluidigm |
| CD197 (CCR7) | 159Tb | G043H7 | Fluidigm |
| CD14 | 160Gd | M5E2 | Fluidigm |
| CD45RA | 170Er | HI100 | Fluidigm |
| HLA-DR | 173Yb | L243 | Fluidigm |
| CD56 | 176Yb | NCAM16.2 | Fluidigm |
| *: CD38 and CD66b labels were changed during the cohort recruitment | | | |

**Table S12. Fluorochrome-conjugated antibodies for membrane antigen staining in Mass Cytometry (CyTOF) analysis.**

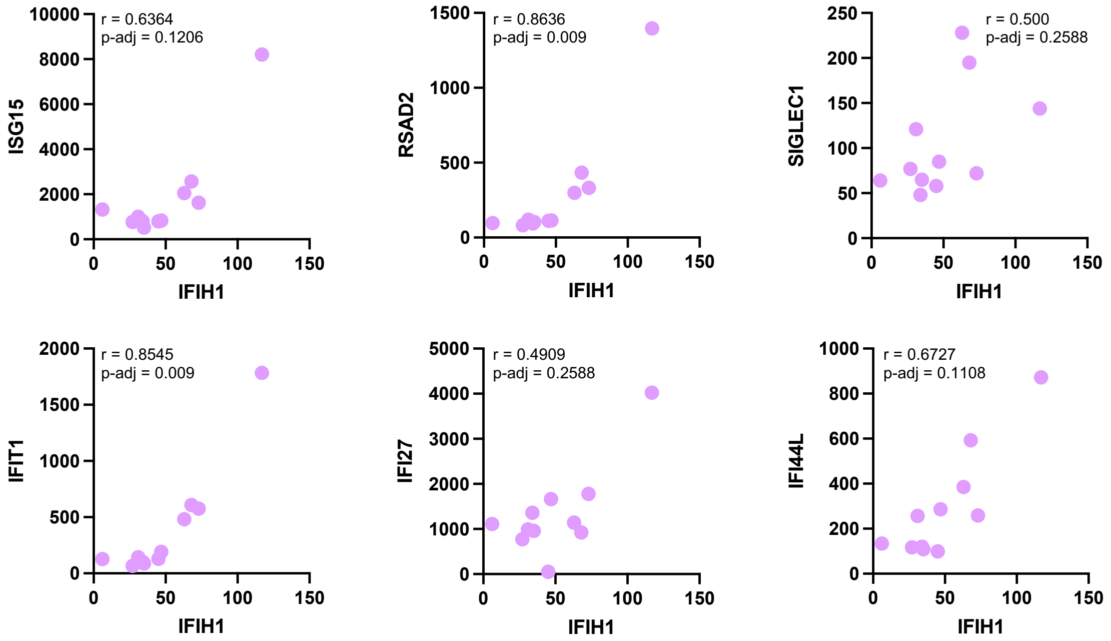


**Figure S1. Correlations between ISGs gene counts and *IFIH1*.** Correlation plot between *IFIH1* gene count and well-described interferon stimulated genes (ISGs) count. Each dot represents a patient. n=11 JDM at diagnosis. p-values are corrected (p-adj) using False Discovery Rate (FDR) method.

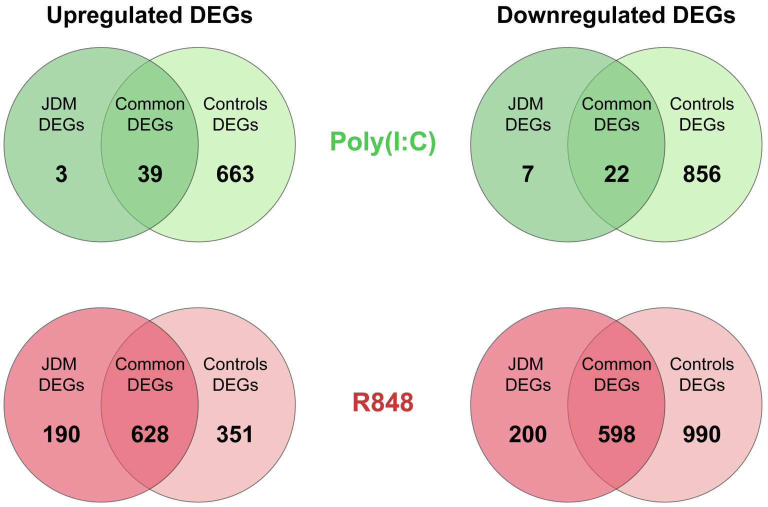


**Figure S2. Number of Differentially Expressed Genes (DEGs) upregulated and downregulated according to stimulation and patients or controls.** Venn diagram representing the number of DEGs after Poly(I:C) (green) or R848 (red) compared to unstimulated condition in patients (JDM) and controls.

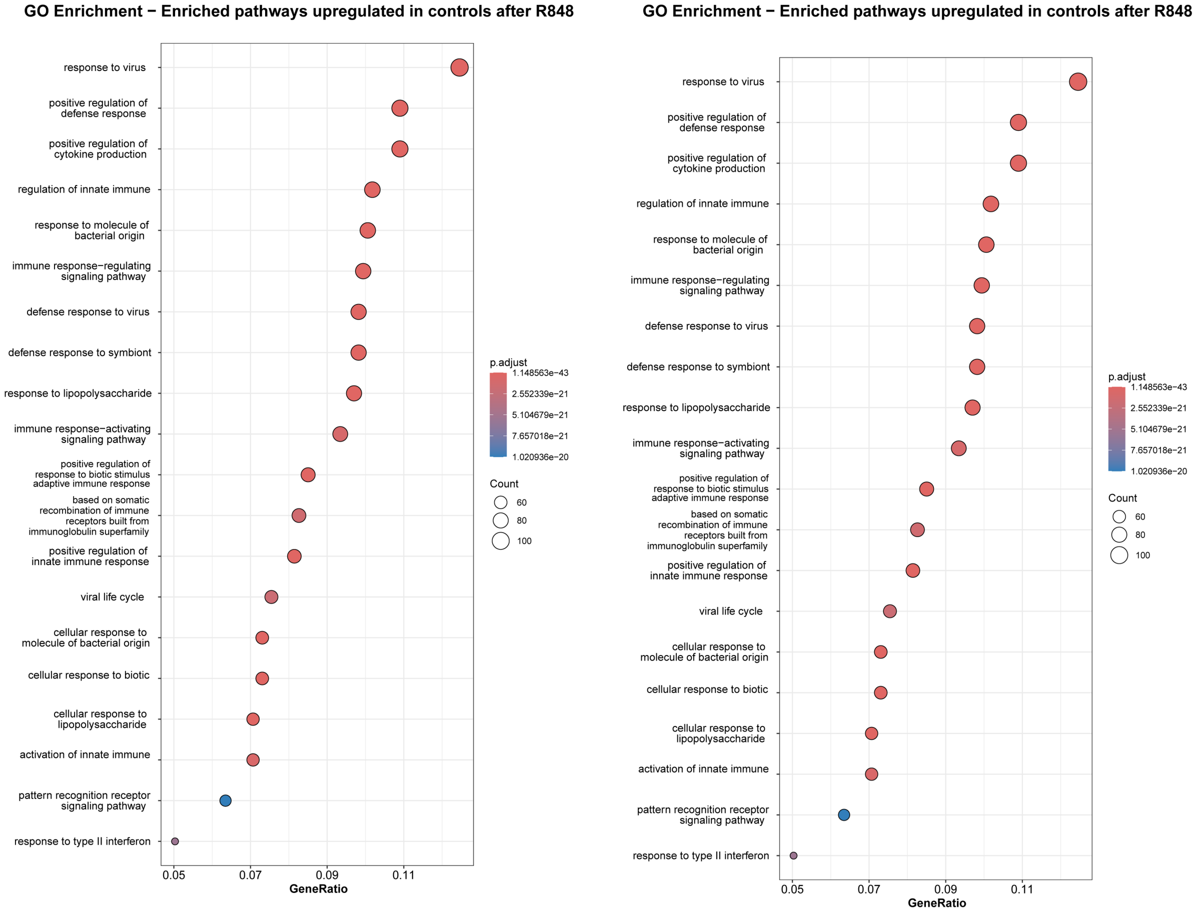


**Figure S3. Enriched pathways upregulated in controls and JDM at diagnosis after R848 stimulations compared to their respective baseline profiles.** Pathways are a list of genes from GO enrichment database. The count represents the number of genes from DEGs that are found in a specific Gene Ontology (GO) term. The GeneRatio is the proportion of our DEGs associated with a particular GO term relative to the total number of our DEGs.

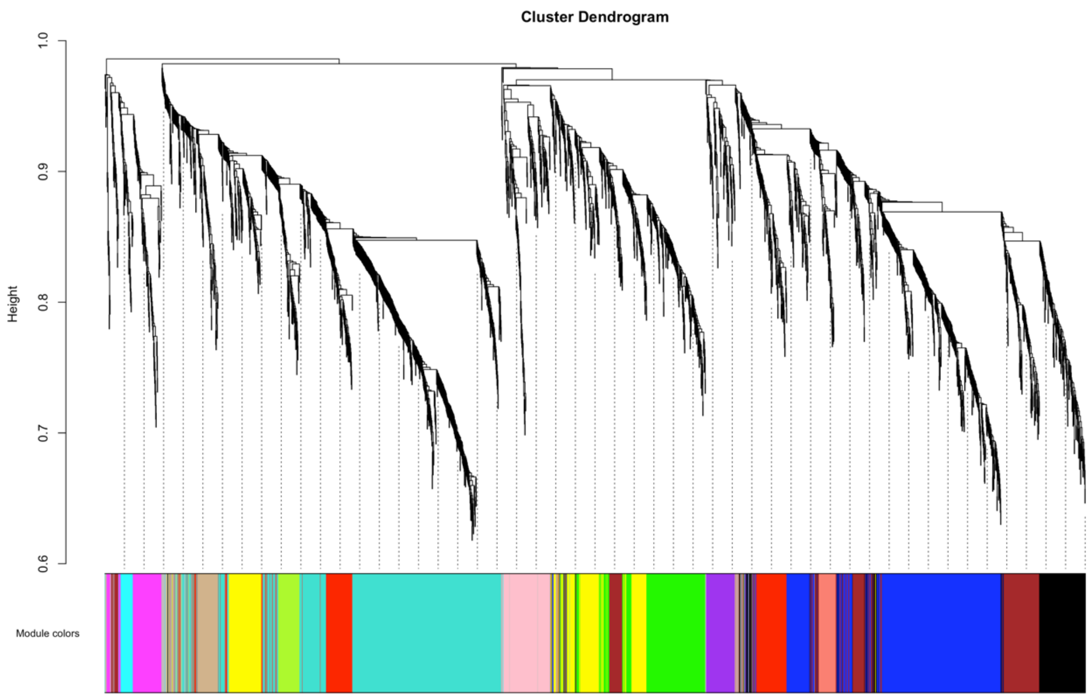


**Figure S4. Dendrogram from Weighted correlation network analysis (WGCNA) reveals distinct genes modules.** This dendrogram shows the hierarchical clustering of genes based on topological overlap in gene expression data, generated using the WGCNA package. Each branch represents a module of color (group of genes with similar expression patterns across the samples). On the y-axis, the height indicates the divergence in gene expression between clusters, with lower values indicating greater similarity.

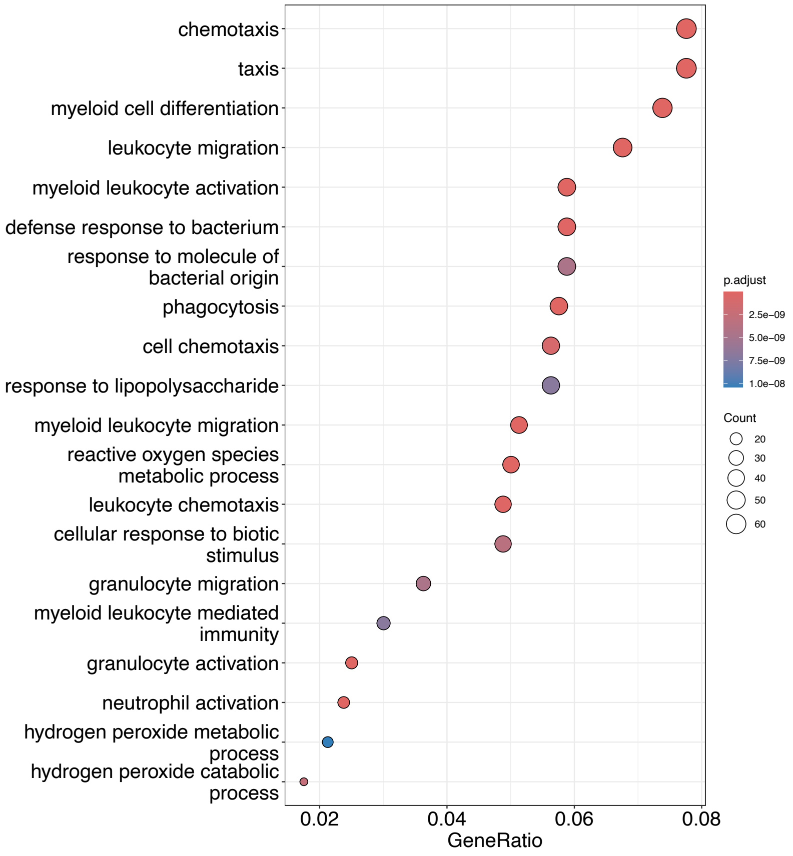


**Figure S5. Enriched pathways analysis from the "turquoise", "greenyellow" and "red gene modules of the WGCNA analysis.** Pathways are a list of genes from GO enrichment database. The count represents the number of genes from DEGs that are found in a specific Gene Ontology (GO) term. The GeneRatio is the proportion of our DEGs associated with a particular GO term relative to the total number of our DEGs.

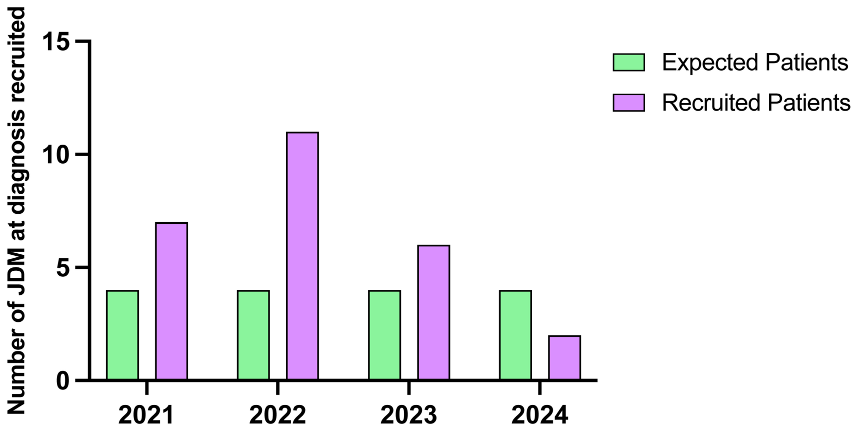


**Figure S6. Number of JDM at diagnosis recruited during the study compared at the number expected before the start of the study based on country incidence.**

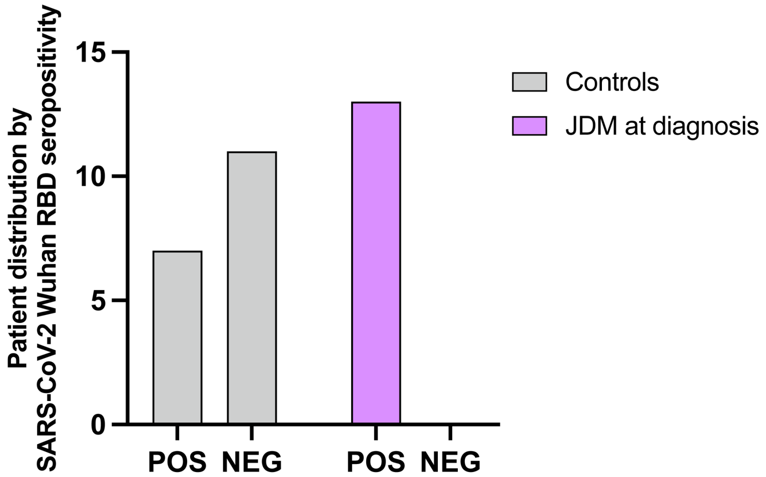


**Figure S7. Histogram showing the distribution of seropositive (POS) and seronegative (NEG) individuals among controls (grey) and JDM patients (purple) for SARS-CoV-2 Wuhan RBD.**
